# Designed Transmembrane Proteins Inhibit the Erythropoietin Receptor in a Custom Binding Topology

**DOI:** 10.1101/2023.02.13.526773

**Authors:** Marco Mravic, Li He, Huong Kratochvil, Hailin Hu, Sarah E. Nick, Weiya Bai, Anne Edwards, Hyunil Jo, Yibing Wu, Daniel DiMaio, William F. DeGrado

**Author notes:** Corresponding Authors Marco Mravic, Daniel DiMaio, William F. DeGrado. These authors contributed equally to this work (Marco Mravic, Li He).

## Abstract

Transmembrane (TM) domains as simple as a single span can perform complex biological functions using entirely lipid-embedded chemical features. Computational design has potential to generate custom tool molecules directly targeting membrane proteins at their functional TM regions. Thus far, designed TM domain-targeting agents have been limited to mimicking binding modes and motifs of natural TM interaction partners. Here, we demonstrate the design of *de novo* TM proteins targeting the erythropoietin receptor (EpoR) TM domain in a custom binding topology competitive with receptor homodimerization. The TM proteins expressed in mammalian cells complex with EpoR and inhibit erythropoietin-induced cell proliferation. *In vitro,* the synthetic TM domain complex outcompetes EpoR homodimerization. Structural characterization reveals that the complex involves the intended amino acids and agrees with our designed molecular model of antiparallel TM helices at 1:1 stoichiometry. Thus, membrane protein TM regions can now be targeted in custom designed topologies.

## Introduction

Protein transmembrane (TM) domains execute diverse and essential biological functions often through molecular features located deep within the bilayer hydrophobic region. It would be advantageous to have chemical biology tools targeting membrane proteins directly at lipid-embedded sites, allowing manipulation of their functional and structural states analogous to how antibodies and small molecules have been applied to water-exposed protein regions^1, 2^. Small molecules like the FDA-approved Eltrombopag, targeting the Thrombopoietin receptor TM region^3^, give precedent for the therapeutic potential of strategies modulating proteins from within lipid.

Approaches for targeting TM regions exist, but remain limited. Truncated peptides mimicking natural TM sequences can perturb assembly of oligomeric complexes and multi-spanning proteins through competition for native inter-TM domain interactions^3, 4–10^. However, the list of membrane protein targets, binding regions, and functional perturbations accessible by simple mimic molecules of known interactions is short. TM polypeptides often have poor solubility and pharmacology. Furthermore, in cell membranes TM polypeptides can exhibit restricted trafficking, insertion with the incorrect topology, and poor binding specificity – limiting their scope as tool molecules.

Rational chemical derivatization^5, 6, 11^, computational design^12^, or expressed TM protein variant libraries^11, 13, 14^ are alternative approaches for TM polypeptide engineering, each offering distinct benefits and limitations. Cell-based phenotypic selections of TM protein libraries have the advantage to yield novel modulators of receptor signaling without knowing the target’s precise structure or essential residues *a priori*^15^. However, functionally selected TM domains often do not form stable complexes with the intended target, and may function through alternative signaling complexes^16^, induced degredation^17^, or transient, indirect, or lipid-specific assemblies difficult to characterize^18^.

Theoretically, structure-based computational design can direct molecules to specific membrane protein regions with custom binding modes, affording the potential to stabilize or recognize distinct protein conformations. However, engineering TM protein complexes has major challenges and sparse precedent. Successful computational design requires both a confident atomic model for the target and accurate estimation of protein interaction energetics in lipid^19^.

Membrane protein force-fields historically have been unreliable, although recent developments are noted^20, 21^. The first *de novo* design proof-of-concept debuted the Computed Helical Anti-Membrane Proteins (CHAMP) algorithm, wherein engineered TM peptides bound and selectively distinguished the highly similar single-pass integrin α_IIb_ and α_V_ TM domains via association of mutual GxxxG motifs presented on both binder and target TM domains^12^, a well-studied high-affinity TM dimerization sequence pattern. Our analogous work a decade later yielded integrin *β*_1_-specific GxxxG-containing TM peptides^22^. Mutagenesis of natural TM domains by computational design to increase affinity of the native oligomer’s existing interaction mode has also been reported: inhibitors of Epstein–Barr virus latent membrane protein-1 assembly via TM5 (67% sequence identity)^23^ and of single-pass BclxL homodimerization regulating apoptosis (87- 91% identity)^24^.

*De novo* computational designs so far have relied on encoding TM association via mimicking a single, known, highly-stable interaction motif. All designs (*de novo* or redesigns) have utilized a single topology: parallel TM helices with type I insertion^23, 25, 26^. Thus, 15 years after the initial proof-of-concept, both the binding modes and spectrum of membrane protein targets have remained extremely limited; computational design of TM-targeting polypeptides is still far from routine.

We sought to develop a computational workflow generally applicable for engineering TM adaptor complexes with customizable geometry. We report several important technological advances in our design of *de novo* TM proteins binding the type I single-span mouse erythropoietin (EPO) receptor (mEpoR) to inhibit signaling. Our primary engineering goal was to encode interaction partners having type II insertion which engage mEpoR in an antiparallel TM helix topology, which is distinct from the native mEpoR homodimer’s parallel topology (Fig. 1a). Thus, our code does not use the receptor’s native TM interactions as a starting point. Secondly, our designed TM mini-proteins can be stably expressed by using a titratable promoter to tune function in mammalian cell membranes, opposed to adding exogenous peptides dosed at saturating concentrations. Finally, we showed that the expressed TM proteins can inhibit a signal-amplifying receptor. EPO binding to only 6% of cell surface mEpoR yields ∼50% maximal cellular proliferation response^27, 28^, imposing a strict functional requirement for our design The designed TM proteins successfully associate with mEpoR and inhibit EPO-EpoR signaling in mammalian cells. *In vivo* and *in vitro* structural characterization define a complex in overall agreement with the guiding design model of a hetero-dimeric antiparallel transmembrane topology. This upgraded CHAMP software implementing RosettaMembrane^29^ can design synthetic TM proteins adopting a specific binding topology with a targeted membrane protein, holding promise for complex tasks of molecular recognition within lipid membranes.

**Figure 1.**
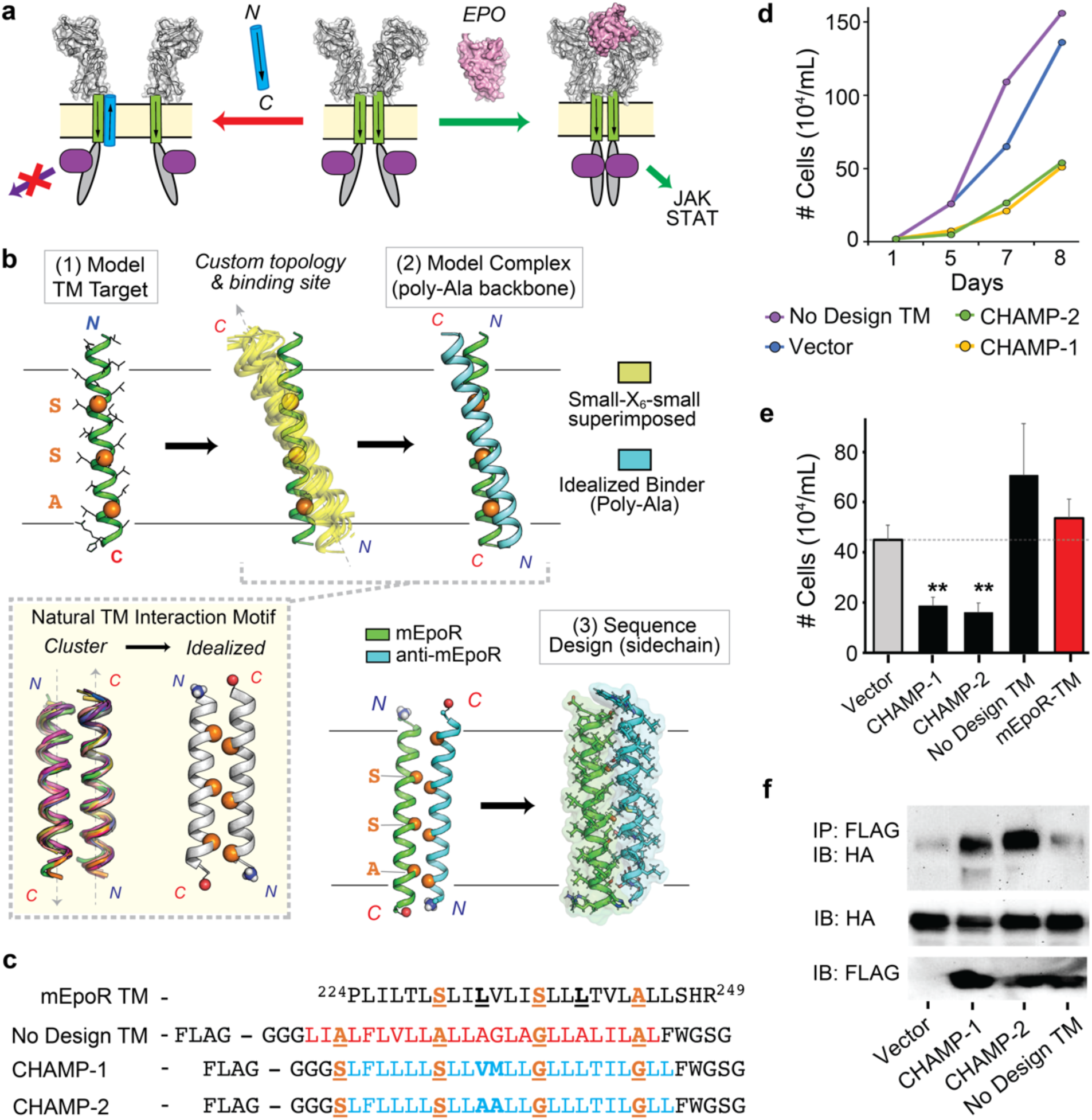
Computational design of anti-mouse erythropoietin receptor (mEpoR) TM domain peptides. (a) Computed Helical Anti-Membrane Proteins (CHAMPs, blue) targeting mEpoR’s TM domain (green rectangle) with antiparallel TM domain topology to competitively inhibit mEpoR’s homodimerization (red arrow) and cross-membrane activation of JAK/STAT signaling (green arrow) induced by EPO (pink, space filling model). Constitutively-bound JAK2, represented in purple. (b) CHAMP Algorithm. mEpoR’s monomeric TM helix (green cartoon) is modeled. Then, a poly-alanine (Poly-Ala) CHAMP binding partner (blue) is positioned to bind mEpoR’s Ser-X_6_-Ser-X_6_-Ala motif (orange, spheres) using a bioinformatics derived template approach, specifying idealized from an antiparallel inter-helical geometry associated with the small-X_6_-small repeat (inset)^38, 39^ extracted from many unique natural membrane protein examples (yellow). Stabilizing sidechain interactions and the *de novo* CHAMP sequence were designed with RosettaMembrane. (c) Top line shows mEpoR TM domain Ser-X_6_-Ser-X_6_-Ala motif (orange) and overlapping Ser-Leu (underlined) repeat. Bottom three lines show expressed FLAG-tagged synthetic TM constructs; No Design control and CHAMP TM regions (red and blue, respectively) and small-X_6_-small motifs (orange, underlined). (d) EPO-induced cell proliferation assay of mouse BaF3/mEpoR cells expressing the empty vector or the synthetic TM domains in IL-3-free media containing 0.06 U/mL EPO. Proliferation is significantly impaired by CHAMP-1 and CHAMP-2 expression. (e) Number of BaF3/mEpoR cells after six days incubation in the presence of 0.06 U/ml EPO (n = 3-10). Error bars, standard error. Asterisks, p-values < 0.05. (f) SDS-PAGE of BaF3/mEpoR cells expressing FLAG-tagged TM proteins immunoblotted either directly (bottom two panels) or after immunoprecipitation (IP) with anti-FLAG antibody (top panel), probed with anti-FLAG detecting TM proteins or anti-HA detecting mEpoR. Representative of 3 experiments.

## Results

### Design of anti-mEpoR TM peptides to bind in a non-native antiparallel complex

EpoR is a prototypical cytokine receptor whose transmembrane domain contributes to both receptor homodimerization^30–32^ and conformational coupling upon ligand binding^33, 34^, activating the Janus kinase (JAK)-signal transducer and activator of transcription (STAT) pathway driving cell proliferation and differentiation. Previously discovered TM domain interaction partners of mEpoR include the Friend retrovirus gp55^35^ and artificial library-derived TM aptamers^15, 18^, which activate EPO-independent signaling through a proposed mechanism of induced dimerization. Mouse EpoR TM domains only weakly homodimerize^30–33^, mediating parallel self-interaction via “Ser-Leu zipper” SxxLxxx 7-residue repeats (Fig. 1a-c)^36^. In model membranes, designed TM peptides hosting SxxLxxx sequence repeats spontaneously homodimerize, strongly favoring parallel TM helix geometry like mEpoR^37^. However, within multi-spanning proteins, SxxLxxx patterns mediate TM helix packing in both parallel and antiparallel geometries^36^. Thus, we hypothesized that mEpoR’s TM domain presents a malleable molecular surface susceptible to target via different geometries: posing the challenge for our algorithm to engage mEpoR via a custom non-native antiparallel topology.

mEpoR’s Ser-Leu zipper also encompasses a 7-residue pattern of small residues every other helix turn: S230-S237-A244. This small-X_6_-small pattern was queried against our published non-redundant membrane protein structural database and globally clustered TM helix-helix interaction geometries^38, 39^, and was found to be associated with common structural topology of tight packing antiparallel TM helices with a shallow left-handed crossing. The consensus small-X_6_-small amino acids directly line these interfaces, allowing close approach to the partner helix’s backbone. From >100 non-redundant natural examples of this 2-helix TM interaction geometry, we generated idealized backbone coordinates: inter-helical distance = 8.1 Å; crossing-angle = -175 degrees, Z-offset = 2 Å (Figs. 1b, S1).

The sequence-structure principles underlying antiparallel interaction of small-X_6_-small TM domains are not well defined, with limited experimental evidence to date – namely case studies of designed repeat TM peptides (e.g. Gly-X_6_-Gly, Ser-X_6_-Ser, Ala-X_6_-Ala)^37, 40^ and EmrE transporter assembly^41^. In contrast with previous design methods, our approach uses data-driven modelling – not relying on receptor mimicry or literature-based “rules” for encoding the desired TM interaction, e.g. GxxxG-mediated motifs^12, 22^. Thus, we test the algorithm’s *de novo* ability to effectively encode a CHAMP sequence by complementary interactions with the target’s unique TM sequence and molecular surface to specify the desired complex’s topology.

First, the target TM domain (here, mEpoR) is modeled as an ideal α-helix embedded in an implicit membrane at an energy-optimized depth and orientation^42^. Second, a poly-alanine backbone model of the putative CHAMP binding partner is built in a favorable helix-helix conformation with the target. The CHAMP was precisely positioned relative to mEpoR’s small-X_6_-small pattern using the aforementioned data-mined idealized antiparallel topology (Fig. 1b). Next, a flexible-backbone sidechain packing routine implemented with RosettaMembrane^43^ designs the CHAMP sequence, optimizing interactions with mEpoR’s TM domain. Of the 24 embedded CHAMP residues, 13 were automatically designated as “potentially interfacial”. The remaining “lipid-facing” residues were semi-randomly assigned a apolar identity (ILVF) fixed throughout design. The 4 small-X_6_-small positions were limited to Gly, Ser, and Ala identities given the bioinformatics data, while the 9 remaining interfacial residues sampled a limited lipid- friendly alphabet (GATSVLIFM). The sequence profile of CHAMP designs shown in Figure S1. The critical final step was ranking and selecting designed sequences by the theoretical stability of their modelled mEpoR-bound complex. Given the documented poor accuracy of predicted interaction energies by RosettaMembrane^19^, we instead ranked models primarily on the lack of sidechain packing voids – a model quality metric commonplace in soluble protein structure prediction and design^44^. We took design models in the top 10% of Rosetta’s “PackStat” score, then reduced to two prominent sequences by sequence clustering (Fig. S1). This complemented rule-based selection of anti-mEpoR CHAMP-1 and -2 sequences, which differed at only 2 interface residues (Fig. 1c). Orthogonal *ab initio* prediction of mEpoR/CHAMP-1 and mEpoR/CHAMP-2 complexes by ESMfold^45^ yielded close-packed models within 0.8-0.9 Å backbone RMSD of our designs (Fig. S1A), suggesting this topology is the lowest energy structure for these sequences.

We also derived a “No Design” TM protein containing a database-derived small-X_6_-small sequence (Fig. 1c; see methods, Fig. S1). This negative control protein allowed testing of binding specificity inherently encoded in a generic small-X_6_-small repeat sequence for mEpoR’s TM domain.

Specific CHAMP algorithm adjustments include (A) implementation in RosettaMembrane to increase user accessibility, (B) ranking designs on interface packing over Rosetta energy scores, and (C) using idealized structural bioinformatics-derived molecular models for the CHAMP/mEpoR complex, versus natural templates. Finally, human visual evaluation was cited as critical in past designs^12^, which introduces user disparities and limits reproducibility. Our adaptations automate model building, design, and final sequence ranked selection – facilitating broader community use.

### CHAMP peptides inhibit EPO-induced growth by binding the mEpoR TM domain

To test the activity of these designed sequences, retroviral transduction was used to stably express FLAG-tagged CHAMP TM proteins in mouse BaF3 cells engineered to express mEpoR (BaF3/mEpoR cells), which lack endogenous EpoR (Fig. 1c-f). Proliferation of BaF3/mEpoR cells can be stimulated by growth factors IL-3 (EpoR-independent) or EPO (EpoR-dependent). In the absence of IL-3 and EPO, CHAMP protein expression does not induce proliferation in the absence of IL-3 (Fig. S2a), indicating a lack of EPO-independent mEpoR activation. As well, IL-3-induced proliferation is not reduced by the designed TM proteins (Fig. S2b), showing that their expression is not cytotoxic.

We next assayed whether the designed TM proteins impair cell proliferation due to EPO-induced EpoR activation. Over 8 days, EPO-treated BaF3/mEpoR cells expressing CHAMP-1 and CHAMP-2 exhibited significantly reduced proliferation with final cell counts reaching 38 ± 5% and 40 ± 6% (average ± S.E.M., n=6), respectively, versus cells transduced with empty vector (p < 0.05). (Fig. 1d-e). EPO-treated BaF3/mEpoR cells expressing the “No design” small-X_6_- small TM protein, a mEpoR TM domain mimic protein, or an unrelated mouse PDGFβR TM domain protein construct^46^ did not inhibit proliferation (Fig. 1e, S2c-d, Table S1). Thus, only the designed CHAMP proteins exerted dominant negative inhibition on mEpoR-dependent proliferation induced by EPO. If the hEpoR was expressed instead of the mEpoR, CHAMP protein expression did not hamper EPO-stimulated proliferation (Fig. S2e). Similarly, CHAMP-1 and CHAMP-2 expression failed to inhibit EPO-stimulated proliferation in cells expressing the “mhm- EpoR” chimera (which consists of the mEpoR’s water-soluble domains but hEpoR’s TM domain, Fig. S2f-g). This specificity for the mEpoR TM domain is remarkable, given that hEpoR differs from the mEpoR by only three mid-spanning residues.

We next used co-immunoprecipitation (co-IP) experiments to test whether the expressed TM proteins physically associate with mEpoR. Detergent lysates of BaF3/mEpoR cells expressing CHAMPs or the No Design control proteins (which are FLAG-tagged at the N-terminus) were immunoprecipitated with an anti-FLAG antibody then immunoblotted with an anti-HA antibody recognizing HA-tagged mEpoR. The FLAG antibody pulled down mEpoR only from cells expressing CHAMP-1 and CHAMP-2 (Fig. 1f), indicated that these proteins formed a stable complex with the mEpoR. Furthermore, the small TM proteins did not affect expression levels of the mEpoR.

Next, we tested the relative association and orientation of mEpoR’s TM domain with the designed TM sequences as cysteine-containing synthetic peptides in model membranes. An established equilibrium thiol-disulfide exchange assay was used, monitoring increased cysteine proximity and reactivity due to formation of non-covalent complexes (Fig. 2a)^47, 48^. Each of the designed TM peptides (C-terminal Cys) were reconstituted with an mEpoR TM domain peptide (N-terminal Cys, mEpoR-TM) in basic buffered solution at 1:100 peptide to detergent or lipid molar ratio. Following glutathione-assisted reversible oxidation, all the disulfide-bonded dimer species were separated and quantified by reverse-phase high-performance liquid chromatography (RP-HPLC). In either dodecylphosphocholine (DPC) micelles or 1-palmitoyl-2-oleoyl-sn-glycero-3-phosphocholine (POPC) small unilamellar vesicles (SUVs), the CHAMP-1, CHAMP- 2, and the No Design Control peptides show a strong non-random preference to form N-to-C disulfide-bonded heterodimers with mEpoR-TM, enriched 2.5 to 16-fold relative to mEpoR homodimers (Fig. 2b-c). Thus, the three *de novo* small-X_6_-small TM peptides form stable antiparallel complexes *in vitro* with mEpoR-TM that outcompete its self-interaction.

**Figure 2.**
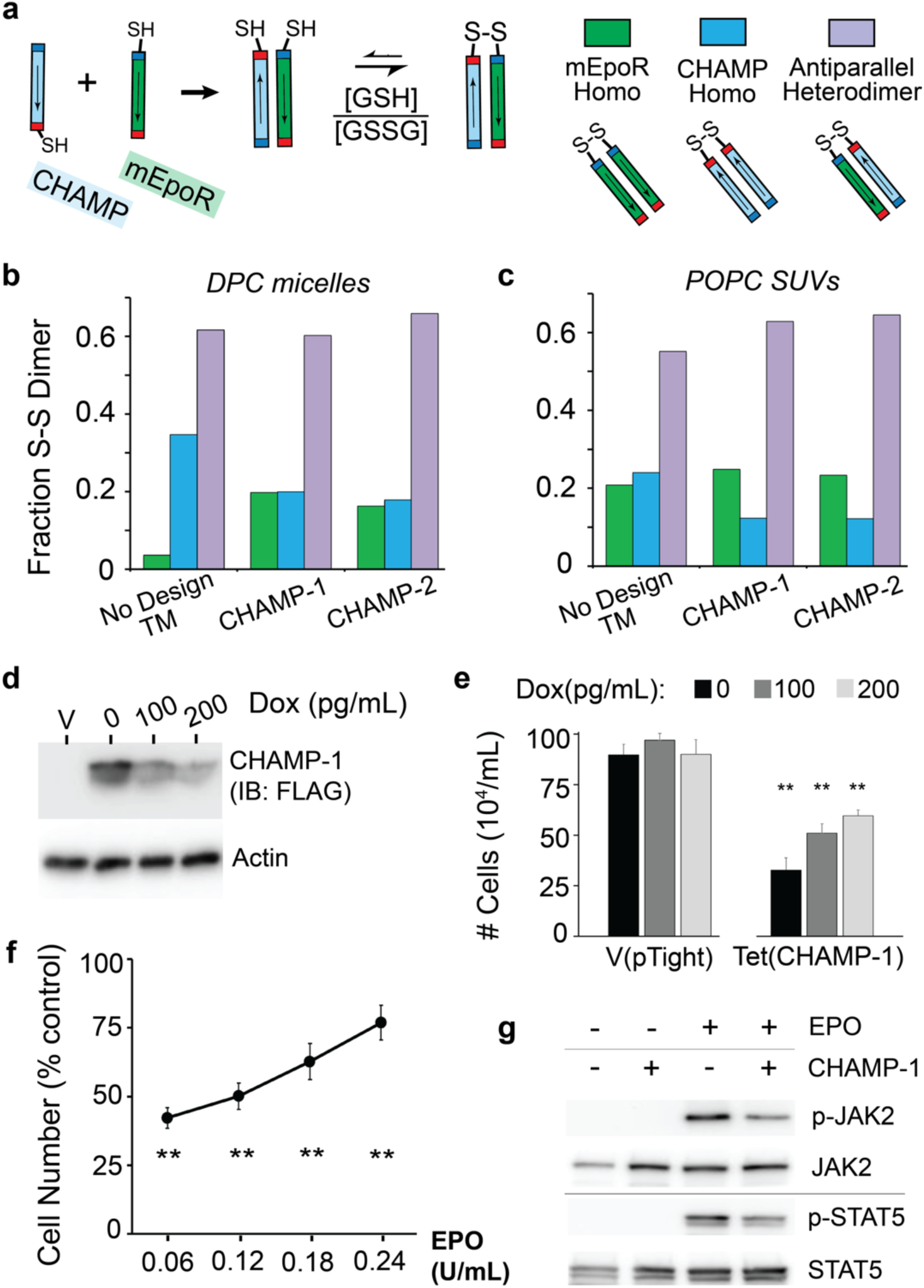
Designed TM peptides target mEpoR TM domain to inhibit the EPO-mEpoR signaling cascade. (a) Equilibrium thiol-disulfide exchange measuring altered thiol proximity due to non-covalent complexes. N-terminal cysteine mEpoR TM peptide (green) and a C-terminal cysteine designed TM peptide (blue) (sequences, Table S2) were solubilized together to 100 µM at 1:50 molar ratio versus model membrane components, (b) dodecylphosphocholine (DPC) detergent micelles or (c) 1-palmitoyl-2-oleoyl-sn-glycero-3-phosphocholine (POPC) small unilamerlar vesicles (SUVs), in 50 mM Tris-HCl pH 8.6, 200 mM KCl, 0.5 mM ethylenediaminetetraacetic acid (EDTA) buffer. Mixtures were reversibly oxidized (10 equivalents glutathione; 3:1 reduced:oxidized). Mole fractions of covalent disulfide-bonded species (S-S dimer) were quantified by reverse phase liquid chromatography: mEpoR homodimer (green), designed heterodimer (purple), designed homodimer (blue). Arrows indicate N-terminal to C-terminal direction. (d) Doxycycline (Dox) repression of FLAG-tagged CHAMP-1 expression levels from a tetracycline-responsive promoter in BaF3/mEpoR cells expressing the tTA tetracycline transactivator, measured by SDS-PAGE and anti- FLAG western blot of cells treated with doxycycline titration. Actin is a loading control. V, empty vector. (e) 0.06 U/mL EPO and either 0, 100, or 200 pg/mL Dox of cells expressing CHAMP-1 or empty V(pTight) vector. (n=3; bars, standard error). (f) Number of BaF3/mEpoR cells expressing CHAMP-1 after incubation for six days in medium containing varying EPO concentration, relative number of cells in the absence of CHAMP-1 expression (n=4). (g) BaF3/mEPOR cell extracts having CHAMP-1 expression and/or 1 U/ml EPO treatment for 10 minutes subjected to SDS-PAGE and immunoblotted with antibodies recognizing either phosphorylated or total JAK2 or STAT5. In panels e and g, statistical significance (p<0.05) versus vector control denoted by asterisks.

### CHAMP-1 inhibits the EpoR signaling pathway in sequence-dependent manner

For CHAMP-1, we investigated the sequence features and mechanism driving its function. Expression under a titratable doxycycline-repressible promoter in BaF3/mEpoR cells showed that inhibition of EPO-induced proliferation is dose-dependent, tunable by CHAMP expression levels (Fig. 2d-e). Likewise, the inhibitory effect was negatively correlated with the concentration of stimulatory EPO (0 to 0.24 U/mL), as expected (Fig. 2f). Phosphorylation-specific immunoblotting showed that EPO-stimulated tyrosine phosphorylation of JAK2 and STAT5, downstream effectors of the EpoR, is reduced by CHAMP-1 expression (Fig. 2g), indicating CHAMP-1 inhibits the EPO-mEpoR cross-membrane signaling axis.

Next, we used mutagenesis to identify amino acids in the mEpoR and CHAMP-1 TM domains required for this inhibition. We first measured the effect of mEpoR mutants containing single and double amino acids substitutions from the hEPOR at the three dissimilar TM positions (Fig. 3a-b). Compared to the 58 ± 5% reduction of final cell count after 6 days upon CHAMP-1 expression in cells expressing wildtype mEpoR (n=10), CHAMP-1 showed similar potency in mEpoR L235V-expressing cells (55 ± 15% reduction) and modestly dampened inhibition in S237L-expressing cells (43 ± 8%). L238V-expressing cells showed significantly reduced responsiveness to CHAMP-1 (20 ± 11% reduction in proliferation; p < 0.003). S237L-L238V- expressing cells were not inhibited (1 ± 13% reduction), while L235V-S237- and L235V-L238V- expressing cells were still partially inhibited by CHAMP-1 (27 ± 4% and 26 ± 8%, respectively). Similarly, a panel of CHAMP-1 mutants was tested (Fig. 3a,c). Mutants S8Q, S8N, S8D, or S8E lose all inhibitory potency; cells proliferate indistinguishably from cells transduced with empty vector. T19Q CHAMP-1 is modestly less inhibitory than CHAMP-1 (30 ± 8% reduction in cell count compared to parental mEpoR/BaF3 cells lacking CHAMP-1, p = 0.01). S1Q was completely tolerated, inducing inhibition similar to wildtype CHAMP-1 (58 ± 7%, p < 0.001). Interestingly, even though CHAMP-1 S8Q failed to inhibit proliferation, this mutant still co-immunoprecipitated mEpoR (Fig. S2h). These mutants could lose potency due to reduced interaction with EpoR or lower monomeric pool of CHAMP-1, given that strongly polar membrane-embedded sidechains often drive TM domain self-association in a depth-dependent manner^49^. CHAMP-1 and CHAMP-2 differ between ^11^V^12^M and ^11^A^12^A, highlighting additional tolerated amino acids. We also explored apolar disruptive mutations. CHAMP-1 small-X_6_-small residues “S1-S8-G15-G22” were mutated to either isoleucine or leucine, I1-I8-I15-I22 (I-I-I-I) and L1-L18-L15-L22 (L-L-L-L) mutants. I-I-I-I CHAMP-1 exhibited significantly lower inhibitory potency, 34 ± 28% reduction in cell number versus the 58% reduction due to CHAMP-1 (p < 0.05), but interestingly did not completely abolish activity (Fig. 3c). By contrast, L-L-L-L lost inhibitory potency and instead induced EPO-independent proliferation similar to previously engineered polyleucine TM proteins (Fig. S2a)^50^, exerting an alternative consequence. We also tested point mutations at four consecutive positions, V11-L14, in an attempt to define the CHAMP-1 helix register binding mEpoR. V11F showed significantly impaired inhibitory activity (24 ± 3% reduction in proliferation, p < 0.05) whereas M12I and L14A exhibited only modestly reduced potency relative to CHAMP-1 not reaching statistical significance (54 ± 9%, 36 ± 16%, respectively). L13A, having the lowest expression level, showed no inhibition (1 ± 3%; Fig. S2i). Thus, mutagenesis did not identify a helix register, as the most impactful V11F and L13A mutations lie on opposite faces of CHAMP-1’s TM helix. Interestingly, *ab initio* predicted^45^ models of each point mutant (Fig. S2j) adopted identical conformations, but revealed mutated sidechains may be tolerated in the interface; for example, V11F adopted a less favorable rotamer. By contrast, the CHAMP-1 I-I-I-I mutant is predicted to be completely non-interacting with mEpoR. Many factors such as mutant expression level (Fig. S2i), membrane trafficking, or reduced monomeric availability may be responsible for apparent reduced potency – factors we did not rigorously quantify. Overall, while many substitutions are tolerated, mutation of the small-X_6_- small motif as well as other sites mitigate CHAMP-1’s ability to inhibit mEpoR.

**Figure 3.**
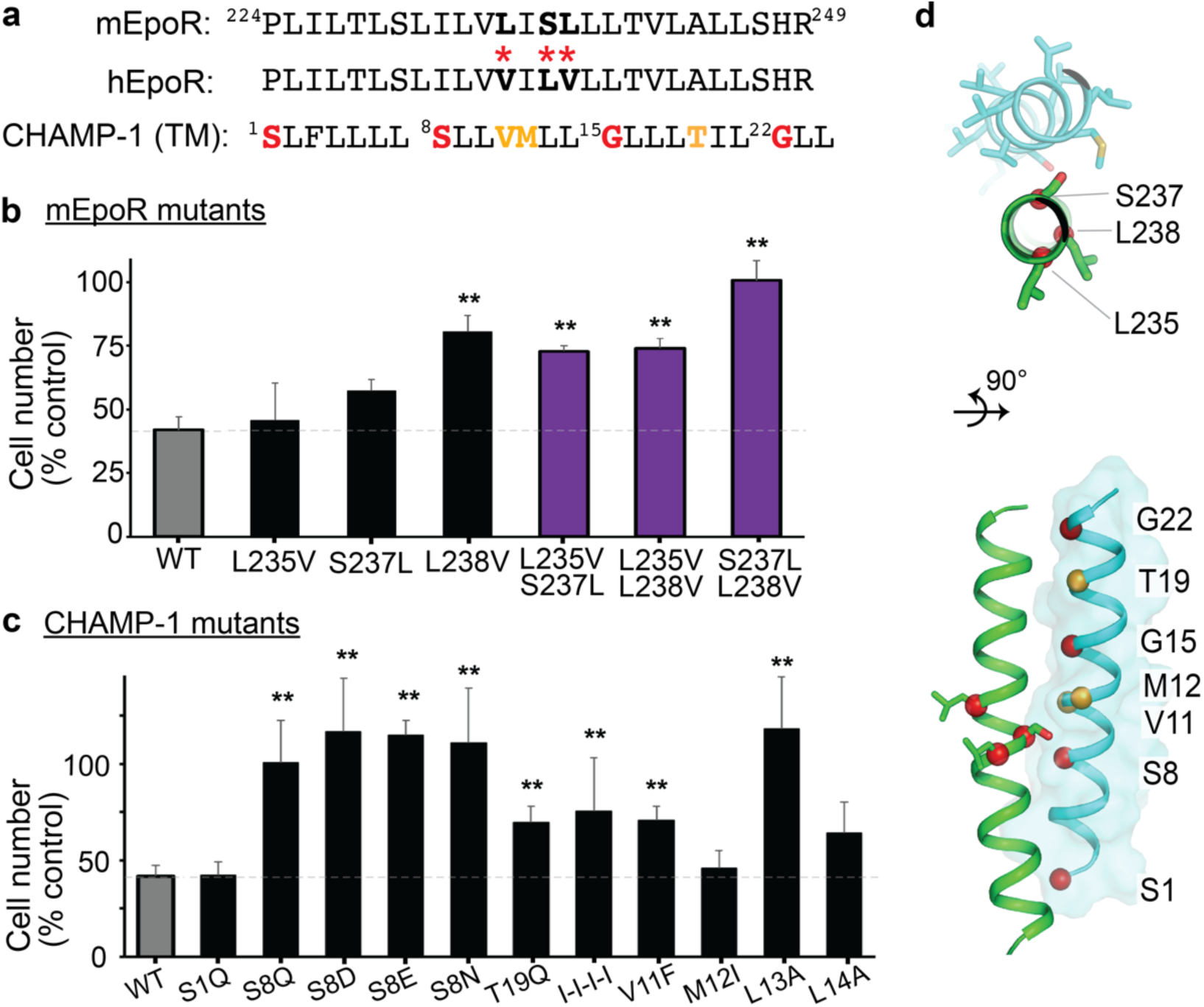
Sequence-specific interaction between mEpoR and CHAMP-1. (a) Wild-type (WT) sequences for core TM regions of mEpoR, hEpoR, and CHAMP-1. Dissimilar residues between mEpoR and hEpoR and indicated by bold letters and red asterisks. Mutated CHAMP-1 residues expected to be in contact with mEpoR from the design model: small-X_6_-small repeat Ser1-Ser8-Gly15-Gly22 shown in red and other amino acids at intermediate helix turns in orange. (b) BaF3 cells stably expressing mEpoR mutants with single or double mEpoR to hEpoR amino acid substitutions co-expressed with WT CHAMP-1 were cultured in medium supplemented with 0.06 U/mL EPO. Day 6 cell counts are shown as a percentage of the number of cells upon EPO-stimulation for BaF3 cells expressing in the absence of CHAMP-1 expression (n = 3; bars, standard error). (c) BaF3/mEpoR cells stable expressing CHAMP-1 mutants were cultured in medium supplemented with 0.06 U/mL EPO. Cell counts at day 6 are shown normalized to the number of cells in the absence of CHAMP-1 expression (n = 3; bars, standard error). Expression levels of some of the mutants were assessed by western blot (Fig. S2). (d) Design model of mEpoR (green) and CHAMP-1 (cyan) TM complex with residues subject to mutation labeled. Top: mEpoR, sticks and red Cα atom spheres; Bottom: CHAMP-1, Cα atom spheres colored as in (a). Significance (p<0.05) in difference versus WT mEpoR with WT CHAMP-1 denoted by asterisks.

### CHAMP-1/mEpoR assemble in anti-parallel orientation in vitro and in mammalian cells

We next structurally characterized the mEpoR and CHAMP-1 TM peptides in detergent micelles solutions. First, we used FRET-based fluorescence quenching to determine the stoichiometry of the complex (Table S2)^51, 52^. Increasing molar ratios of fluorescein-5-maleimide-labeled CHAMP-1 (acceptor) peptide to diethylamino-4-methylcoumarin-3-maleimide-labeled mEpoR (donor) peptide were reconstituted in myristyl sulfobetaine (C14-B) micelles (constant 180:1 detergent to total peptide ratio). Donor emission quenching was observed, decaying linearly to half intensity at 1:1 acceptor:donor ratio (Fig. 4a). Based on theoretical equations (Fig. S3)^52^, the TM peptides appear form a nearly full occupancy complex of 1:1 stoichiometry under these conditions (0.3% mol CHAMP-1). Concentrating the TM peptides by decreasing the detergent:peptide ratio (100:1) did not cause additional quenching (e.g. higher-order assembly), but further diluting complex (250:1) slightly reduced fluorescence decay linearly – increasing the fraction monomer (Fig. S3). These data indicate the fluorophore-labelled CHAMP-1/mEpoR-TM complex is hetero-dimeric as designed. In addition, the interaction affinity *in vitro* is stronger than mEpoR TM peptide homodimerization previous measured in C14B by analytical ultracentrifugation ([monomer] ≈ [dimer] at 0.3% mol)^30^.

**Figure 4.**
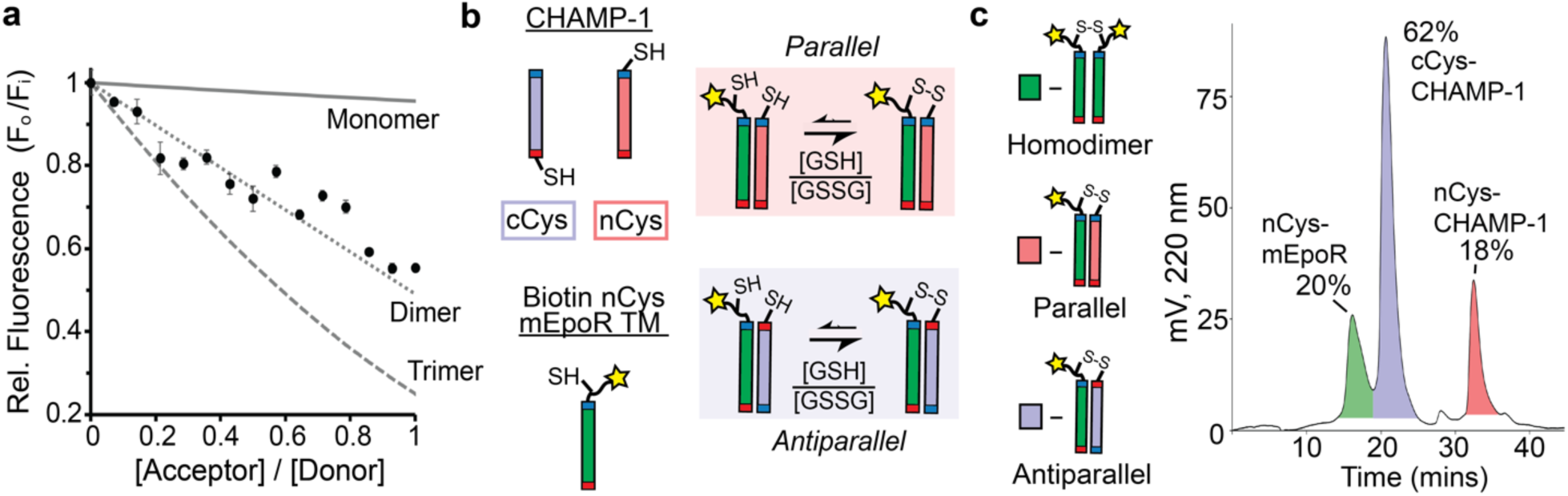
Dimeric antiparallel topology of the mEpoR/CHAMP-1 complex in detergent micelle solutions. (a) Relative (Rel.) 460 nm fluorescence emission of 1.5 µM 7-diethylamino-4-methylcoumerin-labeled mEpoR TM peptide shows FRET quenching as fluorescein-labeled CHAMP-1 is titrated in 0.6 mM C14-betaine, 50 mM Tris- HCl pH 8, 100 mM NaCl, 0.5 mM EDTA, 5 mM TCEP buffer at fixed equimolar total peptide concentration (n=3; bars, standard error). Theoretical FRET curves for monomeric (Mon.), dimeric, and trimeric complexes including a micelle crowding factor (gray). (b) Competitive thiol-disulfide exchange was performed by reconstituting together a biotin-labeled N-terminal cysteine (biotin nCys) mEpoR TM peptide (green) with an N-terminal cysteine CHAMP-1 (nCys-CHAMP-1, red) and C-terminal cysteine CHAMP-1 (cCys-CHAMP-1, purple) at a 1:2:2 molar ratio (0.5 mM total) in 20 mM C14- betaine, then subject to reverse oxidation in buffer from Figure 2a-b (peptide sequences in Table S2). The underlying non-covalent interaction topology of the CHAMP-1/mEpoR complex will strongly influence the propensity of nCys-mEpoR to form disulfide bonds with nCys-CHAMP-1 (parallel interaction) versus cCys- CHAMP-1 (antiparallel interaction). Blue and Red markers on TM domain cartoons represent N-termini and C- termini, respectively. (d) Streptadvidin-bead were used to capture biotin-labelled mEpoR TM containing species: the homodimer (green) and heterodimers with nCys-CHAMP-1 or cCys CHAMP-1. Left, the amount of each monomeric peptide captured by competitive disulfide with biotin-nCys-mEpoR is indicative of the preferred covalent complex formed during reversible oxidation (green, Ncys-mEpoR; pink, nCys-CHAMP-1; purple cCys-CHAMP-1). Right, reverse phase liquid chromatogram of monomeric TM peptides after capture and elution from streptadvidin-beads by reducing agent, injected and separated on a C4 column (Vydac) using an organic solvent gradient of isopropanol/acetonitrile/water/TFA (60:30:9.9:0.1). Representative of n=3 experiments.

We next tested whether the CHAMP-1/mEpoR complex preferentially adopts a parallel or antiparallel orientation through a competitive thiol-disulfide exchange capture experiment (workflow schematic, Fig. S4a). This experiment measures whether a CHAMP-1 peptide containing an N-terminal or C-terminal cysteine (nCys-CHAMP-1, cCys-CHAMP-1) more readily forms disulfide bonds with a biotinylated mEpoR TM peptide containing an N-terminal cysteine (biotin-nCys-mEpoR) in C14B micelle solutions (Fig. 4b, Table S2). A three peptide mixture of 2:2:1 nCys-CHAMP-1:cCys-CHAMP-1:nCys-mEpoR was first reconstituted at 40:1 mole ratio C14B to total peptide, then subject to reversible glutathione-assisted oxidation, followed by low pH quenching (Fig. S4b). mEpoR-containing species were next bound to streptavidin beads, thus capturing TM peptides which had disulfide bonded to mEpoR. After the unbound species were washed off, beads were treated with reducing agent to finally elute all monomeric TM peptides. Species fractions of the collected peptide monomers were quantifying by RP-HPLC (Fig. 4d), revealing a strong preference for cCys-CHAMP-1 at >60% of total peptide captured (>3-fold more than either nCys-CHAMP-1 or biotin-nCys-mEpoR). Thus, the CHAMP-1/mEpoR TM complex predominantly adopts an antiparallel topology *in vitro*.

Finally, we used split green fluorescent protein (GFP) complementation to confirm the antiparallel TM orientation and association of CHAMP-1 with mEpoR in BaF3 cells. In this assay, fluorescence is generated by reconstitution of two non-fluorescent fragments of GFP (GFP1-10 and GFP11) when both fragments are in the same cellular compartment and in the proper proximity and orientation to form a stable complex (Fig. 5a; constructs, Table S1)^53^. The GFP1-10 fragment was fused via a short flexible linker to mEpoR or hEpoR, thereby replacing EpoR’s C-terminal cytoplasmic domain. Control experiments with hEpoR-GFP1-10 or mEpoR-GFP1-10 expressed in BaF3 cells showed low background cellular mean fluorescence intensity (MFI) (2.5×10^3^) (Fig. 5b-c). Next, we co-expressed FLAG-tagged CHAMP-1 with GFP11 fused to the N or C terminus with hEpoR-GFP1-10 or mEpoR-GFP1-10. Co-expression of the N-terminal GFP-11-CHAMP-1 fusion with mEpoR-GFP1-10 led to >4-fold higher MFI than that of cells expressing mEpoR- GFP1-10 alone or the CHAMP-1 fusion alone (Fig. 5b), indicating successful cytoplasmic GFP11 localization and complex formation. We also confirmed complex formation between mEpoR- GFP1-10 and GFP11-CHAMP-1 by co-immunoprecipitation (Fig. S5a). Co-expression of GFP-11-CHAMP-1 with hEpoR-GFP1-10 yielded a smaller 2-fold MFI increase in fluorescence (Fig. 5c), even though mEpoR-GFP1-10 and hEpoR-GFP1-10 are expressed at similar levels (Fig. S5b), consistent with the preference of CHAMP-1 for the mEpoR versus the hEpoR in the growth inhibition assay. In mEpoR-GFP1-10 expressing cells, fluorescence was not increased by expression of alternative non-interacting TM domains (GlycophorinA or ErbB2) fused at their cytoplasmic end to GFP11 (Fig. 5d), although ErbB2 TM domain fusion could complement ErbB2-GFP1-10 as expected (Fig. S5c). The C-terminal CHAMP-1-GFP-11 fusion was not expressed at a detectable level and did not increase fluorescence (Fig. S5a). The ability of CHAMP-1 with N-terminal GFP11 to complement mEpoR with C-terminal GFP1-10 shows that a population of this CHAMP-1 fusion adopts a type II topology with cytoplasmic N-terminus and interacts with mEpoR-GFP1-10 in an antiparallel TM domain orientation, as designed.

**Figure 5.**
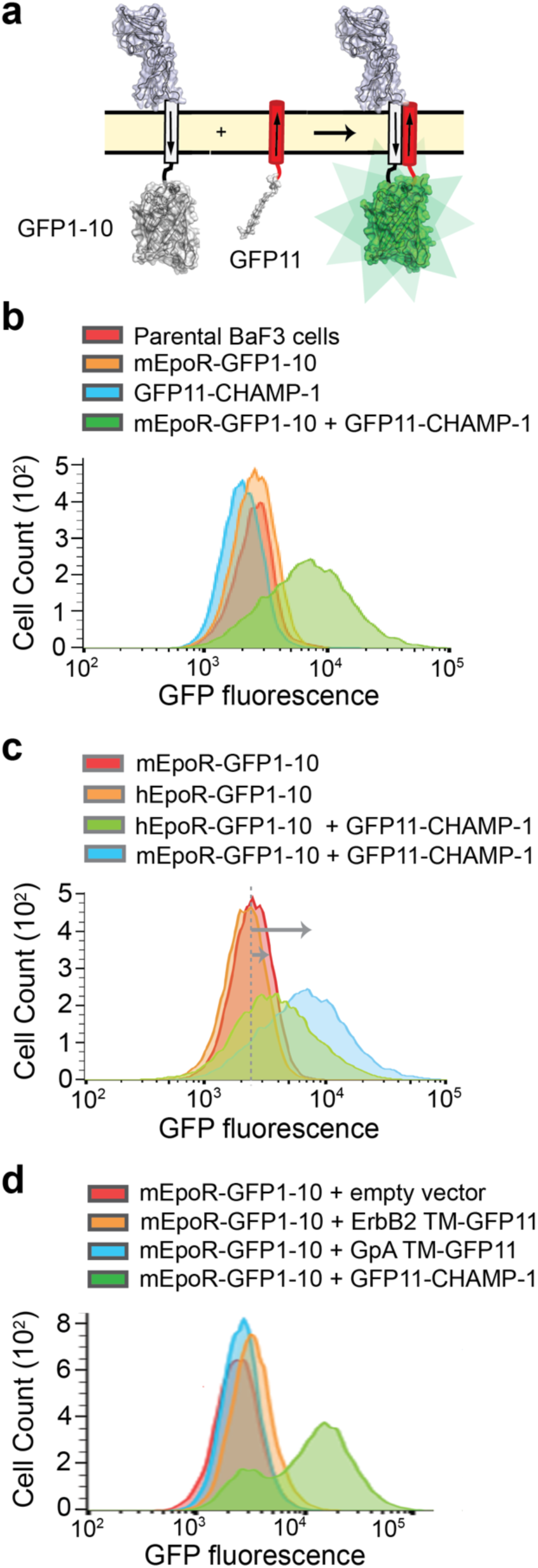
Split GFP assay reveals the anti-parallel topology of mEpoR/CHAMP-1 in mammalian cells. (a) Schematic of split green fluorescent protein (GFP) complementation assay in BaF3 cells. mEpoR and hEpoR constructs (gray) have GFP fragment with β-strands 1-10 (GFP1-10) replacing the C-terminal cytoplasmic domain. CHAMP-1-GFP has an N-terminal fusion to GFP’s 11^th^ β-strand (GFP11), expected to be presented in the cytoplasm, the same cellular compartment as the GFP1-10 fragment if CHAMP-1 adopts type II transmembrane insertion. The arrows represent the N-to-C direction of the TM domains. Fluorescence occurs when GFP1-10 and GFP11 associate. Sequences shown in Table S1. (b) Flow cytometry histogram showing fluorescence of parental BaF3 cells and BaF3 cells expressing mEpoR- GFP1-10 and GFP11-CHAMP-1 together or separately. (c) Flow cytometry histogram showing fluorescence of BaF3 cells expressing mEpoR-GFP1-10 or hEpoR-GFP1-10 in the presence or absence of GFP11-CHAMP-1 (representative experiment, n=3). (d) Flow cytometry histogram showing fluorescence of BaF3/mEpoR-GFP1-10 cells co-expressing empty vector, GFP11-CHAMP-1, or the TM domain of GlycophorinA (GpA) or ErbB2 fused to GFP11.

### Solution NMR shows a direct sidechain-mediated complex at the targeted mEpoR epitope

We next measured solution NMR spectra for isotope-labelled mEpoR TM peptides in the presence of unlabeled CHAMP-1 peptides, repeated for several constructs and membrane mimics (Tables S2, S3). To clearly differentiate chemical perturbations due to CHAMP-1 binding in each situation, we also systematically titrated detergent or bicelle concentration with mEpoR-TM alone to identify spectral changes inherent to its monomer-homodimer equilibrium. First, we recorded ^1^H-^15^N HSQC spectra of [U-^15^N]-labelled mEpoR-TM1 reconstituted with and without 1.3 molar equivalents CHAMP-1 in C14B micelle conditions mimicking our FRET experiment (180:1 detergent:peptide) (800 MHz, 45° C, pH 5.2) (Fig. S6). Distinct new peaks emerged distinct from mEpoR’s homodimer resonances, indicating a slow-exchanging CHAMP-1-bound mEpoR population. A second mEpoR construct (mEpoR-TM2) and different CHAMP-1 peptide having polar TM-flanking sequences were similarly assayed (Table S2). CHAMP-1 titration to U-^15^N mEpoR-TM2 in C14B induced fast-exchanging chemical shift perturbations, while mEpoR-TM2’s monomer-homodimer equilibrium in C14B was in slow-exchange (Fig. S7). In 1,2-dimyristoyl-sn-glycero-3-phosphocholine (DPMC)/1,2-dihexanoyl-sn-glycero-3-phosphocholine (DHPC) q=0.3 bicelles, fast-exchanging chemical shift perturbations were observed for both bicelle and CHAMP-1 titrations yet differed in directionality (Fig. S8), allowing assignment of distinct monomer, homodimeric, and heterodimeric shifts. A dissociation constant of 2.7 ± 0.6 mol % relative to DMPC was fit for CHAMP-1. In DPC micelles, mEpoR-TM2’s well-dispersed monomeric ^1^H-^15^N resonances underwent distinct slow-exchanging behavior upon titration of CHAMP-1 (Fig. 6a-c) and lowered DPC concentration, allowing unambiguous classification of homodimer and heterodimer states (Fig. S9-11). Thus, mEpoR and CHAMP-1 assemble for all model membranes and peptide construct combinations tested, albeit varying in exchange behavior. Surprisingly, these constructs’ association affinities inferred from population fractions were much weaker than expected from our FRET experiments, possibly attributed to chemical changes at peptides’ termini or lower pH NMR buffer. Nonetheless, the CHAMP-1/mEpoR-TM2 complex was isolated as the major species in the sample with spectra suitable for resonance assignment. DPC gave the best intensities, linewidths, and dispersion when both detergent (>400:1 DPC:mEpoR) and CHAMP-1 (>1.5 mol %, or 6-8 equivalents) were in excess.

**Figure 6.**
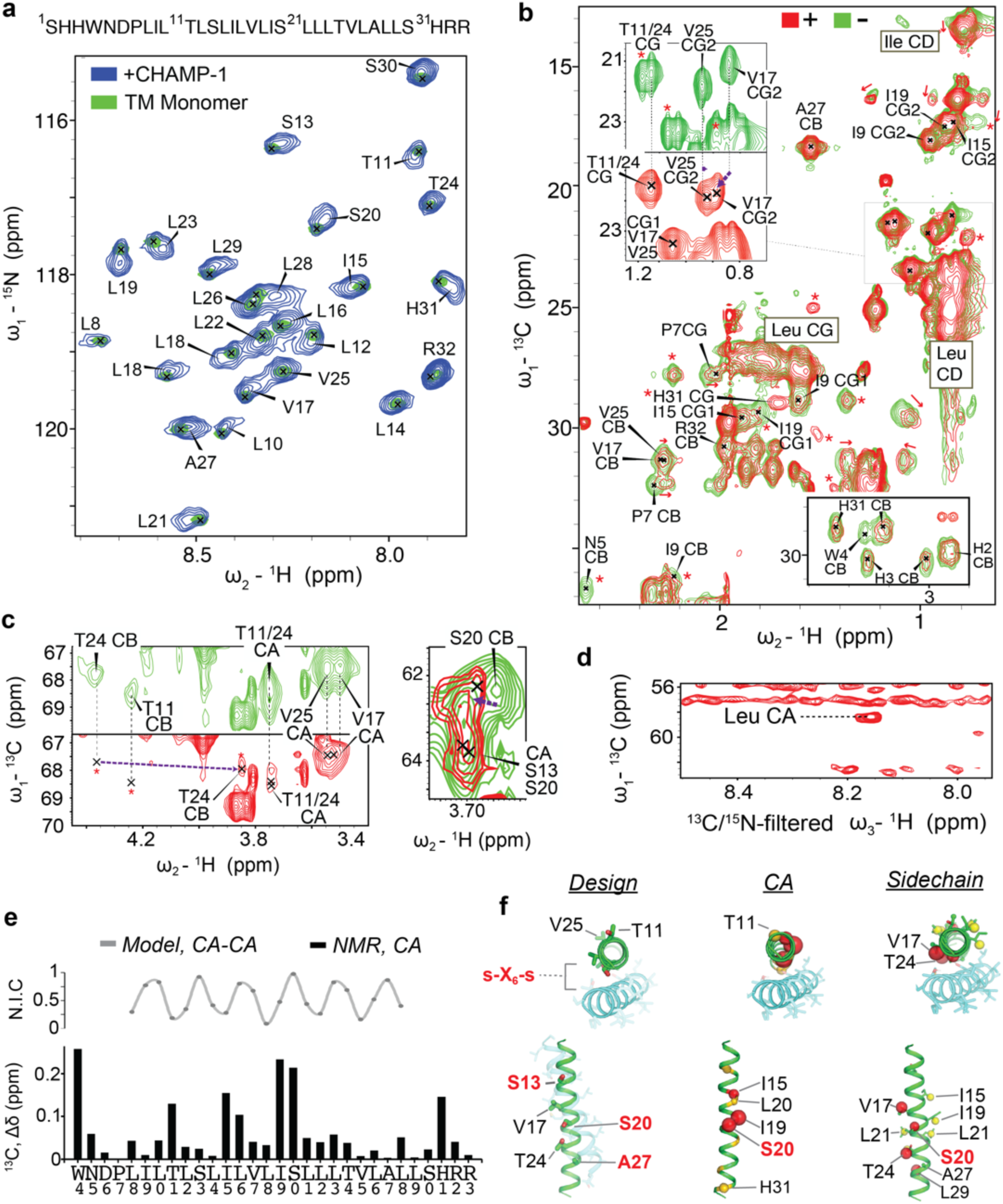
Solution NMR of the side-chain mediated CHAMP-1/mEpoR complex in DPC micelles. (a) mEpoR-TM2 sequence and ^1^H-^15^N HSQC spectra in ^2^H-DPC at 200 µM (U-^15^N^13^C^1^H) with 40 mM sodium acetate pH 5.2, 20 mM NaCl, 0.5 mM EDTA, 5 mM dithiothritol (45° C, 800 MHz). Monomeric (green; peaks, X’s) and CHAMP-1-bound states (blue, 1 mol %) independently assigned. (b) ^1^H-^13^C HSQC spectra of mEpoR-TM2 monomer (green) and CHAMP-1-bound (red) states from (a) have widespread differences: chemical shift perturbations (red arrows); new or broadened peaks (red asterisks). *Top inset* (5x contour), V17 CG2-HG2 peak shift. (c) Target epitope residues. *Left*, shift perturbation of T24 CB (purple arrow), new unassigned peaks (red asterisks), broadening of T11 CA/CB and T24 CA resonances. *Right*, shift perturbation of S20 CB-HB resonance, alongside broadening of S13 CA, S13 CB, and S20 CA. (d) 2D F1-^13^C-edited / F3-^13^C,^15^N-filtered HSQC-NOESY spectrum. Transferred NOE cross-peaks indicate direct contact between mEpoR-TM2 ^13^C atoms, e.g. Leu CA, and CHAMP-1 ^14^N/^12^C-attached protons, e.g. backbone amide proton(s). (e) Helical periodicity in mEpoR-TM2 CA shift perturbation upon CHAMP-1 binding and its agreement with expected CHAMP-1/mEpoR CA-CA distances from the design model, plotted as normalized inter-helical closeness (N.I.C., see methods). (f) *Left*, CHAMP-1 (cyan) mEpoR (green) design model noting protein- and lipid-facing sidechains (sticks) and targeted small-X_6_-small repeat (red). *Middle*, perturbed CA atoms (scaled spheres, green-red color scale) lie on one face of mEpoR’s TM α-helix, overlapping the targeted epitope and CHAMP-1 interface; minimally perturbed CA are lipid-facing in the design model. *Right*, perturbed sidechain atoms enriched at one helix face, including V17, S20, T24.

In ^2^H-DPC, we characterized how CHAMP-1 binding alters mEpoR-TM2’s sidechain environment. We first assigned [U-^15^N, ^13^C]-labelled mEpoR-TM2’s monomeric state in excess ^2^H-DPC using triple resonance HNCA and HNCB spectra followed by (H)CC(CO)NH and H(CC)(CO)NH backbone-sidechain TOCSY spectra to separate heavily overlapping ^1^H-^13^C spectral regions (Fig. S9, S12-13). Then, we performed independent backbone and partial sidechain assignment of the mEpoR-TM2/CHAMP-1 resonances (Fig. S14). Comparison of monomeric versus CHAMP-1-bound mEpoR-TM2 ^1^H-^13^C HSQC spectra showed induced broadening of select peaks, new resonances, and widespread chemical shift perturbations across diverse sidechain chemical groups (Fig. 6b-c, S15). Numerous mEpoR sidechains in close interaction (< 4 Å) with CHAMP-1 in our design model experienced significant changes, including V17 CG2 (shift), I19 CG1 (broadening) and CG2 (shift), T24 CB (shift), Ser13 and Ser20 overlapping CA’s (broadening), and Ser20 CB (shift). Induced ^1^H-^13^C chemical shift perturbations upon CHAMP-1 binding was similar to that of homodimeric mEpoR in about 50% of resonances, including numerous assigned (e.g. V17, A27, S30, H31, R32) and unassigned peaks (Fig. S15).

Likewise, CHAMP-1 induced many shift and intensity perturbations distinct from mEpoR homodimerization, including at mEpoR-TM2 residues contacting CHAMP-1 in the design model (I9, I19, S20, T24) and membrane-proximal residues (H3, W4, N5, P7).

Interestingly, per-residue shift perturbations (Δ*δ*) to mEpoR-TM2’s ^13^CA atoms exhibited a clear pattern of 3-4 residue periodicity (Fig. 6d), mirrored to a lesser extent by ^1^H-^15^N Δ*δ* (Fig. S16), possibly indicating its helical register and interaction surface with CHAMP-1. Notably, Δ*δ* is roughly in-phase with the inter-helical CA-CA distances between mEpoR and CHAMP-1 within the core of our design model (Fig. 6e-f). mEpoR-TM2 CA atoms with the largest Δ*δ* are closest to CHAMP-1, while residues least perturbed are lipid-facing. However, this correlation was not exact and became out of phase towards both helix termini, suggesting a slightly different CHAMP- 1 packing angle. Thorough comparison of sidechain resonances along the mEpoR-TM2 TM helix was obfuscated by spectral overlaps, particularly for leucines. All the unambiguously assigned resonances with significant spectral changes are plotted on mEpoR’s structure in Figure 6f. While CHAMP-1 binding has a dispersed impact, a greater quantity of highly perturbed sidechain atoms lie at the helix face expected to bind CHAMP-1 (e.g. V17, I19, S20, T24).

Finally, we measured direct inter-atomic interaction between ^1^H,^13^C,^15^N-mEpoR-TM2 and unlabeled CHAMP-1 in ^2^H-DPC through a F1-^13^C-edited/F3-^13^C,^15^N-filtered HSQC-NOESY experiment. Intermolecular transfer resonances were observed between ^13^C-attached protons (mEpoR) and ^12^C- or ^14^N-attached protons (CHAMP-1) in the 2D projection spectrum (Fig. 6d). One cross-peak can be attributed to a CHAMP-1 amide proton (8.14 ppm) interacting with an mEpoR-TM2 leucine ^13^CA (58.4 ppm). Overlap of many leucine CA peaks prevented unambiguous residue-specific assignment (Fig. S15). Nonetheless, these results indicate the complex between CHAMP-1 and mEpoR-TM2 in DPC is direct and features tight backbone-backbone packing as in the small-X_6_-small-mediated design.

## Discussion

Here, we develop and successfully demonstrate a more automated and distinct implementation of the CHAMP algorithm wherein membrane protein-specific bioinformatics data guide the design of *de novo* TM domains to bind a target protein with conformational specificity. To assess whether our procedure indeed encoded the intended custom binding mode with the mEpoR TM domain, we undertook rigorous biophysical characterization typically absent in TM design studies to date. The CHAMP-1/mEpoR TM complex forms readily and is stable *in vitro* across diverse membrane mimics, elevated temperatures, and pH, attributed to an extensive interaction network. CHAMP-1’s robust sidechain-mediated binding contrasts that of past library-selected synthetic EpoR-activating TM domains which co-immunoprecipitated with EpoR, but direct complexes were not detected in detergent given negligible sidechain chemical shift perturbations to EpoR TM fragments measured by solution NMR^16, 18^. Furthermore, our stably expressed mini-membrane proteins exhibit exquisite molecular recognition in lipid bilayers, discriminating between the highly similar hEpoR and mEpoR TM domains. Fluorescence quenching, thiol-disulfide exchange, and split GFP experiments of the CHAMP-1/mEpoR complex *in vitro* and *in vivo* are consistent with the intended non-native 1:1 heterodimeric antiparallel TM topology designed *in silico*.

The precise structure and sidechains stabilizing the complex are not fully clear and consistent between our cellular and biophysical data. NMR spectra in DPC micelles show residues I236 and S237 of the mEpoR are most strongly perturbed (i.e. I19, S20 of mEpoR-TM2 peptide) along with V234 (V17) and T241 (T24). These residues constitute a continuous helix face shared with the mEpoR TM surface (S230-S237-A244) we targeted in our design, implying their interaction with CHAMP-1 (Fig. 6f). Yet, CHAMP-1’s inhibition in BaF3 cells was most sensitive to mEpoR mutations at L238 and S237, but not I236, suggesting the former residues contact CHAMP-1. Notably, both sets of experiments are consistent with S237 participating in the complex. However, the TM helix register implied by interface mapping using activity measurements in living cells differ between solution NMR perturbations in DPC. Residues L238 and I236 lie on opposite faces of mEpoR’s TM domain (∼200° helix rotation apart) thus cannot both simultaneously contact monomeric CHAMP-1. It is possible different conformations of TM helix packing and ensembles of interacting amino acids exist in each distinct chemical environment. There is precedent for similar behavior, with integrin *β*_3_ having alternative TM interfaces inferred from experiments performed in human versus bacterial cell membranes^54^.

On CHAMP-1, substitutions at small-X_6_-small positions reduced or reversed CHAMP-1 influence on BaF3/mEpoR cell proliferation, indicating small-X_6_-small residues are important for interaction. V11F, in phase with the small-X_6_-small motif, slightly reduced inhibition. Yet, the L13A mutation on the opposite helix face fully abrogated inhibition, and mutations at adjacent sites (L14A, M12I, and V11A-M12A within CHAMP-2) had minimal effect. Interestingly, the negative control design assembles into antiparallel complexes with mEpoR-TM *in vitro,* suggesting the small-X_6_-small motif is sufficient to encode interaction. However, this TM protein control did not co-IP with mEpoR or inhibit signaling in mEpoR/BaF3 cells, indicating this sequence is missing features critical for *in vivo* interaction and activity in the more heterogeneous cellular environment. Conversely, S8Q shows no inhibitory activity, but still immunoprecipitates mEpoR. Thus, the exact structure-activity relationship of our designed TM domains remains difficult to parse out from inhibitory activity *in vivo*. This is not entirely surprising, as many important membrane protein properties that affect anti-EpoR activity in cells can be altered upon TM protein mutation: expression level, cell surface trafficking, additional TM interactions competitive with mEpoR binding (either self-assembly or off-target), etc. Nonetheless, the intended mechanism of CHAMP-1 was successfully encoded, namely binding competitively to the mEpoR TM domain in an antiparallel orientation leading to mEpoR signaling inhibition.

We demonstrate design of small expressible synthetic TM proteins completely from scratch with target TM domain specificity and control of interaction topology to effectively perturb a signal-amplifying surface receptor. The CHAMP-1 sequence may serve as a novel tool to complement the existing suite of engineered water-soluble polypetides and TM agents for studying the EPO receptor’s signal conduction mechanism^55–58^ and role in erythropoiesis and other activities^59^. The technological advances described here should facilitate accessibility and increasing complexity in design of tool molecules that target diverse membrane proteins directly at their bioactive TM regions.

## Methods

### Computational Design

We first built an all-atom 25-amino acid model of the mEpoR monomeric TM domain in its lowest energy orientation in an implicit bilayer using RosettaMembrane FastRelax protocol implemented in RosettaScripts^43^ with score function weights: mpframework_smooth_fa_2012^38^. We next built a model of a second interacting TM domain helix (CHAMP) guided by our previous database of the most common geometric motifs between interacting pairs of TM domain helices present in natural membrane protein X-ray structures^38^. Of the >1000 examples of helix-helix interactions extracted from membrane protein PDBs that clustered into the top 15 most prevalent repeated geometries, 141 natural TM helix-helix examples of unique sequence fell into the cluster whose consensus sequence is a repeat of small amino acids every 7 residues (small-X_6_-small). The geometry of this cluster was close-packing anti-parallel TM helices with a left-handed crossing. Of these, we selected those examples whose sequences contains three consecutive small-X_6_-small motifs on each interacting TM helix and being at least 22 residue per helix. Six TM helix pairs matched these criteria. For these six structures, we calculate best-fit coiled-coiled parameters of their backbone coordinates to Crick’s coiled-coil equations for the special case of C2 anti-parallel symmetry. With parameters representative of these structure’s helix-helix geometries (listed in main text), we generated *de novo* coordinates for the corresponding idealized dimeric anti-parallel coiled-coil using Crick’s parametric equations – implemented using CCCP octave source code describe in reference ^60^. One helix of the idealized anti-parallel coiled-coil model was aligned with the mEpoR TM domain model in register with its expected small-X_6_-small motif (heptad position ‘a’), thus creating a knowledge-based template for the positioning a second poly-alanine CHAMP helix for close packing with the adjacent mEpoR small-X_6_-small motif. This process of templating diverges from the past two CHAMP design algorithms, in that past work templated the position of a new CHAMP using exact coordinate of single examples of Small-xxx-Small-motif containing natural examples of TM domain helix pairs extracted from membrane proteins. These examples often contain non-ideal helix geometries. Here, we use ideal geometries in a single coordinate set derived from structural informatics data of the most relevant TM helix-helix templates matching the sequence pattern and TM domain complex we intended to stabilize.

Using this poly-alanine model, we performed sequence design of the CHAMP sequence using rotamer trials in RosettaMembrane, albeit with a very limited sequence alphabet guided by our informatics data. The designation of each sequence position (interface, lipid-facing) and the sequence logo output are shown in Figure S1c. Interface positions at the small-X_6_-small motif were limited to A, S, or G. Interface positions at alternate helix turns were limited to A, S, T, V, I, M, L, F. Lipid-facing residues were allowed to be selected from this alphabet during the RosettaModeling, but were later replaced by selecting residues randomly as previously described^22^: A, I, V, F at 10 % probability and L at 60 % probability, with no additional Ala-X_6_- Ala or Ala-X_3_-Ala motifs being allowed to form at lipid-facing positions. Design was performed using Rosetta LayerDesign, first packing sidechain rotamer trials at core positions, then interface boundary positions, and finally non-interface position. One round of these three rotamer trial steps was followed by a FastRelax step, which includes Cartesian minimization, rotamer repacking (with fixed sidechain identity), and rigid body re-orientation of the heterodimeric TM domain complex in the implicit bilayer to minimize its insertion/solvation energy. Each model was evaluated for the absence of large packing voids at the helix-helix interface by RosettaHoles^44^, i.e. the PackStat filter, and its total Rosetta energy.

The summary of 2000 independent protein design trajectories via these two metrics is plotted in Figure S1c. Due to the documented poor performance of this then-state-of-the-art potential function for computing accurate interaction energies for membrane proteins^19^ and its lack of correlation with the PackStat metric (Fig. S1b), we chose to rank models predominantly based off their PackStat score. The top-ranking models were those in the top 10% of PackStat score (>0.62) and greater than the average value of Rosetta total energy. The resulting sequences and their variability are represented in the weblogo plot in Figure S1c. We next clustered these sequences using the BLOSOM85 matrix and hierarchical clustering. The BLOSOM85 matrix was justified because the calculated median pairwise identity between the top subset of desgined sequences was ∼85%. Three sequence clusters resulted. One CHAMP cluster differed from the other two by having Ser rather than Gly at the final small-X_6_-small position, which induced non-ideal helix geometry at the neighboring mEpoR TM domain. Thus, this cluster of sequences was discarded. The sequences with the top PackStat score from each the remaining two clusters are the core TM domain interface residues of CHAMP-1 and CHAMP-2, which only differ at two mid-spanning residues that lie on the intermediate helix turn between small-X_6_-small residues (Fig. 1). The final molecular model of CHAMP-1 and CHAMP-2 had in common two separate inter- helical hydrogen bond networks involving mEpoR’s Ser and Thr sidechains, displayed in Figure S1a. All steps were performed using scrips that automated the decision-making process using the above described rules for model building, design, ranking, and sequence selection. The only human intervention and rational decision-making in the design algorithm was choosing to target the small-X_6_-small with the relevant TM domain motif and choosing to reject one of the top-ranked sequence clusters.

A “No Design” TM domain sequence was selected whose interface residues are directly derived from a natural small-X_6_-small sequence example of two interacting TM helices extracted from a natural membrane protein, detailed visually in Figure S1d-g. We sought to select a sequence as a negative control peptide and a marker for how productive protein design was at dictating protein-protein interaction specificity and occupancy in binding mEpoR within cell membranes (as opposed to self-interaction or alternate interactions). We began with the same 6 template models from the anti-parallel TM structural motif cluster containing mutually interacting small-X_6_-small sequence. We searched through the 12 TM domain sequences from the 6 pairs of extracted TM domain templates for a TM domain sequence where if mEpoR’s sequence is threaded on to the interaction partner (aligned to the small-X_6_-small motif) there are no significant unavoidable sidechain steric clashes in the resulting model. Fixed identity repacking rotamer trials was performed using the Rosetta fixed backbone (‘fixbb’) application. Only in 1 of the 12 cases was an acceptable model found where no steric clashes were present. It is important to note that the amino acids of the eventual “No Design” control TM domain that faced mEpoR’s TM domain in this model were not altered and the sidechain packing was specifically selected for being optimized towards mEpoR’s molecular surface (Fig. S1f) - just for the lack of major clashes.

The source structure and TM domain antiparallel helix pair sequence of “No Design” control TM was Photosystem II light harvesting complex (PDB:3bz1, chain B), comprised of TM domains 1 and 2, where TM helix 1 contributed the interface residue of the No Design TM. TM helix 2, on which mEpoR was threaded, only has 22% sequence identity with mEpoR (Fig. S1e), which is essentially random given the statistical distribution of amino acids in membrane proteins. TM helix 1’s interface residues were selected (Fig. S1g) while its lipid-facing residues were replaced with the same semi-random selection of apolar residues as in the CHAMP design. Replacing these residues were important given the number of polar amino acids present in this TM domain which are likely impactful for Photosystem II stability and function, but would be unfavorable as a potential mEpoR binding molecule. We additionally added 2 residues to the C-terminal end of the TM domain, Ala and Leu, to match CHAMP sequence designs in the number of small-X_6_-small residues, at 4, but ensuring that residue is not the last apolar residue. The final sequence selection and comparison of the “No Design” control TM relative to the Photosystem II source TM domain sequence is shown in Figure S1f, at 48% sequence identity.

Constructs for protein expression were designed such that the N-terminus and FLAG tag could be in the cytoplasm and a neutral C-terminus was long enough past the TM domain that the charged carboxylic acid is in the lumen. Synthetic peptides of CHAMP-1 were designed specific to each experiment, varying the composition of polar residues flanking the TM domain, but usually included a Trp residue for spectroscopic detection.

ESMfold^45^ server was used to predict *ab initio* the lowest energy structures of mEpoR and CHAMP complexes. Restraints from the lipid bilayer (geometry or insertion) are not considered in this prediction, i.e., the prediction is similar for water-soluble proteins. The input sequence included the mEpoR TM domain span shown in Figure 1 connected at the C-terminus to a 10- residue poly-glycine linker (to allow conformational flexibility) followed by the CHAMP sequence. Predictions performed with a 20-glycine linker gave the same structures. Thus, a parallel helix orientation is a possible outcome but was not returned by the algorithm

### Peptide synthesis and purification

TM peptides were synthesized by solid-state synthesis using fmoc chemistry on rink amide resin in microwave-heated reaction vessels by robotic liquid handling (Biotage Initiatior+ Alstra), cleaved using a trifluoroacetic acid (TFA) cocktail (Sigma) from solid-phase resin, and purified by reverse phase HPLC as previously described^22^. All peptides were produced as C-terminal carboxamides and with free amino N-termini, except for biotin-nCys-mEpoR. Precusor nCys- mEpoR peptide was labeled at its free amino N-terminus as a protected peptide on resin with NHS-Biotin (Sigma) by swelling the resin with DMF having 10 equivalents N,N-Diisopropylethylamine, then adding 1.5 mol equivalents of NHS-Biotin dissolved in minimal DMF to be stirred at room temperature for 45 minutes – performed twice. Peptides were purified by RP-HPLC using a C4 prep column (10 µm 214TP Vydac) using a linear gradient of solvent A (water, 0.1% TFA) and solvent B (60/30/9.9/0.1 isoproanol/acetonitrile/water/TFA). Peptide purity of >95% was achieved in all cases, confirmed using analytical HPLC (C4 Vydac). Correct product masses were confirmed by MALDI mass spectrometry using the matrix α-cyano-4- hydroxycinnamic acid (Sigma).

Peptides for fluorescence quenching cCys CHAMP-1 and nCys mEpoR were derivatized in solution with fluorescein-5-maleimide and (diethylamino-4-methylcoumarin-3-yl)maleimide, respectively (Anatrace). Four mg lyophilized peptide (TFA salts) was dissolved with 10 molar equivalents maleimido-fluorphore in 1 mL DMF and 0.2 mL water with pH 7 HEPES (4-(2- hydroxyethyl)-1-piperazineethanesulfonic acid) buffer (final, 25 mM) was added followed by incubating the reaction overnight at room temperature under nitrogen gas on a rotating shaker. Product fluorescent labelled peptides were re-purified by HPLC as described above.

### FRET-based Fluorescence Quenching of TM peptides for Stoichiometry Determination

Following previous published protocols^51^, we reconstituted donor (diethylamino-4- methylcoumarin-3-yl)maleimido) labeled mEpoR TM peptide at different donor to CHAMP-1 acceptor molar ratios yet at a fixed total peptide concentration across the titration, fixed peptide:detergent ratio, and total equimolar mEpoR to CHAMP-1 ratio. This was done by co-dissolving mEpoR (from a stock solution in trifluoroethanol, TFE, stock solution) with a mixture of unlabeled CHAMP-1 and fluorescein-5-maleimido CHAMP-1 (separately as ethanol stock stock solutions) and C14-B (TFE stock solution) at final concentrations of 1.5 µM donor-labeled mEpoR and 1.5 µM CHAMP-1 (combined unlabeled and acceptor-labelled) with C14-B at 0.5 mM (∼100:1 detergent:peptide), 0.725 mM (∼175:1), or 0.95 mM (∼250:1); assuming that C14- B,CMC = 0.2 mM and has 90 detergent monomers per micelle. The peptide/detergent mixtures were evaporated, then reconstituted in 50 mM Tris-HCl pH 8, 100 mM NaCl, 0.5 mM EDTA, 5 mM TCEP, subject to bath sonication (30 minutes), vortexed (3 minutes), equilibrated overnight, then aliquoted in triplicate into 96-well black round-bottom plates and read in a SpectraMax H5 fluorescence monochromator (Molecular Devices). Fluorescence emission scans were recorded after excitation at 410 nm (435 nm cut-off) and relative intensity at 460 nm was plotted for samples of increasing acceptor-labelled CHAMP-1 mole ratios ([donor]/[acceptor]). Emission intensity decay was compared to theoretical quenching expected for different donor/acceptor molecular complexes different stoichiometry using classic theoretical equations (see ref ^52, 61^). To these theoretical equations, we also added a probabilistic micelle crowding factor using Poisson statistics to calculate the mole fraction of donor additionally quenched due to the probability of non-specific co-occupation of the same micelle as a acceptor peptide, given the concentration of C14-B in the micelle phase, C14-B’s aggregation number, and the detergent:peptide ratio.

### Thiol-disulfide Equilibrium Exchange

For the two peptide mixtures (Fig. 2a-b) samples were prepared by mixing TFE stock solution of nCys mEpoR, ethanol stock solution of cCys designed TM peptide, and methanol stock solution of either DPC or POPC to a final total peptide to detergent ratio of 1:100 and equimolar peptide ratio. Solutions were dried under nitrogen gas and stored under vacuum overnight, then reconstituted at 100 µM concentration of each peptide in 100mM Tris–HCl pH 8.6, 100 mM KCl and 1 mM EDTA with 0.45 mM oxidized (GSSG) and 1.05 mM reduced (GSH) glutathione to initiate reversible redox conditions. After equilibrium was reached overnight, samples were quenched by acidification by adding HCl to 0.1 M final concentration. Each reaction mixture was separate by analytical reverse phase HPLC using a C4 column (Vydac 214TP, 5 µm) and eluted peaks were collected and identified by mass spectrometry. Peaks from the UV chromatogram were integrated to quantify the relative species fractions of disulfide-bonded dimer species: homodimers and heterodimers.

To determine the preferred interaction topology (parallel versus anti-paralel) a novel biotin capture thiol-disulfide exchange procedure was performed to isolate only the mEpoR-containing species from the 9 possible monomeric or disulfide-bonded species when 3 cysteine-containing peptides (Figure 4b-c, Table S2) are mixed together for competitive reversible oxidation. Samples were prepared by co-dissolving peptides and detergent in organic solvent, evaporating solvent, and reconstitution in aqueous solution. Biotinylated nCys-mEpoR was reconstituted at 100 µM with 4-fold molar excess of CHAMP, 200 µM nCys CHAMP-1 and 200 µM cCys CHAMP-1, in 20 mM C14-B (40:1 detergent to total peptide ratio) in 20 mM Tris pH 8, 50 mM NaCl as well as 3 mM glutathione (5:1, [GSH] / [GSSG]), allowed to undergo reversible oxidation overnight. SDS-PAGE confirmed TM peptide oxidation (Fig. S4). The mixture was quenched by reducing the pH to 4.5 by addition of concentrated sodium acetate and bound in batch to streptavidin-conjugated biotin beads overnight – capturing a fraction of the biotin-nCys-mEpoR monomeric and disulfide bonded species. Non-covalently associations were diluted and excess unbound TM peptides were removed by washing the beads (5 bead volumes, 5 times) with 200 mM C14-B, 100 mM sodium acetate, 50 mM NaCl. Peptides disulfide bonded to mEpoR were eluted from the beads by washing with 10 mM TCEP added to the same high C14-B content buffer, reducing disulfide bonds and also diluting non-covalent TM peptide interactions. The eluted fractions were pooled, concentrated and separated by analytical RP-HPLC using linear solvent gradient with C4 column to quantify relative species mole fractions as described above.

### Recombinant Expression of Isotopically Enriched mEpoR TM Domain Fragments

Constructs encoding the mEpoR TM domain fused to His-tagged T4 lysozyme (cysteine-free mutant) having a thrombin cleavage site or to His-tagged SUMO having a sequence-specific nickel-assisted cleavage (SNAC) site were cloned into a pET28a(+) vector (Table S3). Proteins were expressed in BL21(DE3) or C43 cells in M9 minimal media supplemented with 0.5 g ^15^N NH_4_Cl_2_ (Cambridge isotopes) and/or 3 g ^13^C glucose (Sigma) per liter as well as 0.2 g/L of isotope-enriched ISOGROW supplements (Sigma). Cells were induced with 0.4 mM IPTG at an optical density (600 nm) of 0.8 followed by overnight growth at 30 °C or 37 °C. Cells were pelleted, subjected to freeze thaw, and reconstituted in a lysis buffer effective at solubilizing inclusion bodies: 8 M urea, 0.5 mM EDTA, 50 mM Na Phosphate pH 7.5, 2% (w/v) sodium dodecyl sulfate (SDS). Cycles of tip sonication (10 minutes) and rotary shaking (30 minutes) were repeated until a clear homogeneous (non-viscous) solution was achieved. After centrifugation to remove any insoluble debris (30 minutes, 35,000 g), the lysate was poured over a gravity column of Ni-NTA agarose resin (HisPur, Thermo Fisher). The Ni-NTA column was washed with 10 column volumes of detergent-free lysis buffer, then washed with 4 column volumes of 25 mM imidazole detergent-free lysis buffer, before elution in lysis buffer containing 1% SDS and 250 mM imidazole. For thrombin cleavage, the eluted fraction was concentrated and buffer exchanged into 50 mM Tris pH 8, 100 mM NaCl with 0.1% n-Dodecyl-B-D-Maltoside to remove excess SDS by using an 30 kDa amicon centrifugal filter (1/10 dilution, 3 times; EMD Millipore) before a final overnight dialysis (1/500 dilution) using a 20 kDa membrane (Slide-a-lyzer, Thermo). For SNAC cleavage, imidazole was removed (<0.5 mM) using an amicon 10 kDa centrifugal filter without any excess SDS removal steps and buffer exchanged to 100 mM N-Cyclohexyl-2-aminoethanesulfonic acid (CHES) pH 8.5, 100 mM NaCl. The SNAC peptide self-cleavage reaction was initiated by addition of NiCl_2_ to 2 mM final concentration. Due to the high residual SDS content we allowed in this purification (reducing the rate of the SNAC reaction below the typical >95% cleavage in 16 hours), the SUMO-SNAC-mEpoR-TM2 construct underwent SNAC cleavage at 42 °C and typically reached >95% completion in 24-36 hours. The cleaved isotope-enriched mEpoR TM domain peptide was then purified by RP-HPLC and lyophilized, typically yielding ∼10 mg TM peptide per 1 L culture.

### Solution NMR in membrane mimics

Samples were prepared from ethanol or methanol stocks of known concentration of isotopically-enriched recombinant mEpoR TM domain fragments and synthetic CHAMP-1 peptides, combined with lipid or detergent, dried under nitrogen gas stream, and further dried under vacuum. Samples were reconstituted NMR buffer, 40 mM sodium acetate pH 5.2, 20 mM NaCl, 0.5 mM EDTA, 10 mM DTT, 5 % (v/v) D_2_O. The samples were filtered (0.2 µm) and transferred to a 3 mm Shigemi tube. ^1^H-^15^N and ^1^H-^13^C HSQC spectra of labeled mEpoR fragment were recorded at 45° C on a Bruker 800 MHz spectrometer with cryogenic triple-resonance probes, either titrating the concentration of unlabeled CHAMP peptide added or varying the model membrane mimic composition and concentration. Peptide and detergent concentrations of each sample are detailed in each figure legend. For the monomeric mEpoRTM-2 samples, HNCA and HNCB as well as (H)CC(CO)NH and H(CC)(CO)NH backbone-sidechain TOCSY were recorded on the same Bruker 800 MHz spectrometer for a sample of 400 mM ^2^H-DPC and 1 mM ^1^H,^15^N,^13^C mEpoR-TM2. For the CHAMP-1 bound mEpoR sample, HNCA, HNCB, ^13^C-edited NOESY-HSQC, and ^13^C-edited/^13^C,^15^N-filtered HSQC-NOSEY were recorded for a sample of 400 mM ^2^H- DPC, 1 mM ^1^H,^15^N,^13^C mEpoR-TM2, and 7 mM unlabeled CHAMP-1 on a Bruker 800 MHz spectrometer. As well, HNCA and HN(CO)CA were recorded for a sample comprised of 800 mM ^2^H-DPC, 2 mM ^1^H,^15^N,^13^C mEpoR-TM2, and 12 mM unlabeled CHAMP-1 on a Bruker 900 MHz spectrometer with a triple resonance cryogenic probe.

For mEpoR in 10% (w/v) DMPC/DHPC q=0.3 bicelles titrated with CHAMP-1, the nearly saturating chemical shift perturbations were fit by non-linear least squares to a biomolecular interaction scheme using each peptide’s concentration as its mole % (100 × mole fraction) relative to DMPC:

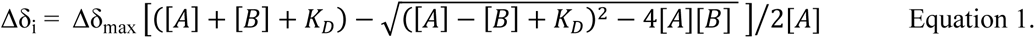

where [A] is mEpoR’s concentration and [B] is the CHAMP-1 concentration expressed in mole %, while K_D_ is the bimolecular dissociation constant, Δδ_max_ is the maximum chemical shift perturbation observed, and Δδ_i_ is the chemical shift at the particular CHAMP-1 concentration measured. K_D_ was globally fit across all titration series fit while each Δδ_max_ was locally fit for each individual resonance titration series.

Chemical shift perturbation at CA atoms were normalized (0 to 1) to the largest induced shift value and plotted on the monomeric TM helix of mEpoR (Fig. 6e), thusly scaling the relative sphere size and color (green, least perturbed; red, most perturbed) of each CA atom. For sidechain atoms, atom’s ^1^H or ^13^C having measured shift perturbation were split into three groups by the magnitude of this perturbation (green, least perturbed; yellow, modestly perturbed; red, most perturbed), not include resonances significantly broadened beyond detection (S13, T11).

### Cloning and Vectors for Mammalian Cell Expression

The HA-tagged hEPOR and HA-tagged mEPOR genes were originally obtained from S. Constantinescu, Ludwig Institute, and subcloned into pMSCV-neo (Clontech) using EcoRI and HpaI restriction sites. The chimeric mhmEPOR and mEPOR mutants containing point mutations in the mEPOR TMD were constructed using double-stranded DNA gBlock^TM^ Gene Fragments (Integrated DNA Technologies), as previously described^18^. The construct encoding the human PDGF-β receptor TM domain flanked by a signal sequence was described previously^46^. The hEPOR-GFP1-10 fusion protein was constructed by replacing the C-terminus of hEPOR downstream of residue 258 with a 10-amino-acid flexible linker (GGSGGGGSGG) followed by the sequences encoding the GFP 1-10 fragment (residue 1 – 215) using DNA gBlock^TM^ Gene Fragments and BglII restriction sites. The mEPOR-GFP1-10 fusion protein was constructed by replacing the sequence encoding hEPOR1-258 in hEPOR-GFP1-10 with the sequence encoding mEPOR1-257 using DNA gBlock^TM^ Gene Fragments and HpaI/BstBI restriction sites. Similarly, GFP1-10 was fused after the TM domain of ErbB2. All non-inducible GFP11 fusion proteins were constructed by cloning DNA gBlock^TM^ Gene Fragments into pMSCV-neo by using EcoRI and XhoI restriction sites. The doxycycline-responsive ErbB2TM-GFP11 was constructed by cloning DNA gBlock^TM^ Gene Fragments into pTight-puro by using BamHI and EcoRI restriction sites. The general structure of the ErbB2 and GlycophorinA TM-GFP11 proteins is hEpoR signal peptide – TM domain – GFP11.

### Cells, retrovirus infections, and growth inhibition assays

Human embryonic kidney (HEK) 293T cells were maintained in DMEM-10 medium: Dulbecco’s Modified Eagle’s Medium (DMEM) supplemented with 10% fetal bovine serum (FBS) (Gemini Bioproducts), 4 mM L-glutamine, 20 mM HEPES (pH 7.3), and 1X penicillin/streptomycin (P-S). To produce retrovirus stocks, 2 µg pantropic pVSV-G (Clontech), 3 µg pCL- (Imgenex), and 5 µg of the retroviral expression plasmid of interest were mixed with 250 µl of 2x HEBS. 250 µl 0.25 M calcium chloride was then added with bubbling into each mixture. The mixture (∼500 µl) was incubated for 20 minutes at room temperature and then added drop- wise into 2.0 x 10^6^ 293T cells plated the day before in 100 mm tissue culture dishes in DMEM-10. The cells were incubated with the transfection mixture for 6-8 hours at 37°C, and the medium was replaced with 5 mL fresh DMEM-10 medium. The cells were incubated for another 48 hours at 37°C, then the viral supernatant was harvested, filtered through a 0.45 µm filter (Millipore), and either used immediately or stored at -80°C.

Murine interleukin-3 (IL-3)-dependent BaF3 and derivative cells were maintained in RPMI-10 media: RPMI-1640 supplemented with 10% heat-inactivated FBS, 5% WEHI-3B cell-conditioned medium (as the source of IL-3), 4 mM L-glutamine, 0.06 mM β-mercaptoethanol, and 1X P-S. BaF3 cells expressing mEPOR, hEPOR, and all EPOR mutants and chimeras were generated by infecting BaF3 cells with pMSCV-neo vector containing the desired HA-tagged EPOR gene. 5×10^5^ BaF3 cells were washed with phosphate buffered saline (PBS) and then re-suspended in 500 µl RPMI-10 medium with 4 µg/mL polybrene. 500 µl retroviral supernatant or 500 µl DMEM-10 for mock-infection was added to the re-suspended cells, which were then incubated for eight hours at 37°C. After incubation, 9 ml RPMI-10 was added and the cells were incubated overnight at 37°C prior to selection in 1 mg/mL G418. Wild-type and mutant CHAMP proteins cloned in MSCV-puro were introduced into cells by infection, followed by selection in 1 µg/ml puromycin.

For proliferation assays, 2×10^5^ BaF3 and derivative cells expressing the appropriate genes were washed in PBS three times to remove IL-3. Cell pellets were resuspended in 10 mL RPMI- 10 medium lacking WEHI-3B cell-conditioned medium but supplemented with 0.06 U/ml human erythropoietin (Epoetin Alfa, Amgen). Viable cells were counted six to eight days after IL-3 removal. All growth inhibition assays were performed in at least three independent biological replicates (*i.e.,* independent infections to express TM proteins). All reported experiments included positive and negative controls that performed as expected, and no outliers in these experiments were excluded. All graphs show average cell counts +/- standard error of the mean (SEM). Statistical significance of differences between control and experimental samples was evaluated by either one-tailed or two-tailed Student’s t-tests with unequal variance, performed using T.TEST function in Microsoft Excel (2013).

### Construction and analysis of inducible cell lines

BaF3 cells were transduced to express an engineered version of the tetracycline-controlled transactivator protein, tTA-Advance (tTA), via retroviral infection with the pRetroX-Tet-Off Advanced (Clontech) vector and selection with 1 mg/ml G418. CHAMP-1 or ErbB2TM cloned in the expression vector pRetroX-TIGHT-puro (Clontech) was introduced into cells expressing tTA by retroviral infection and selection with 1 µg/ml puromycin. HA-mEPOR was retrovirally transduced with pMSCVneo (Clontech) and selected with 0.6 U/ml human erythropoietin (Epoetin Alfa, Amgen) in the absence of IL-3.

To assess expression levels of the CHAMP proteins, BaF3/mEPOR/tTA cells expressing a CHAMP protein were grown in 10 ml cultures in RPMI-10/IL-3 medium in the absence of doxycycline (DOX) or supplemented with 100 or 200 pg/ml DOX for 48 hours. Cells were pelleted in the presence of 1mM phenylmethylsulfonyl fluoride (PMSF) for 10 min at 1,500 RPM at 4°C. Cell extracts were prepared and 20-30 µg of total protein was electrophoresed. After transfer to 0.2 micron PVDF membranes and blocking in 5% milk in TBST (20 mM Tris, 150 mM NaCl, 0.1% Tween 20), blots were incubated overnight 4°C with 1:1000 Anti-FLAG-hrp (Sigma-Aldrich) in 5% Milk in TBST. Blots were then washed and visualized using enhanced chemiluminescence.

For proliferation assays, BaF3/mEPOR/tTA cells expressing CHAMP proteins were first cultured in 10 mL RPMI-10/IL-3 medium in the absence of DOX or supplemented with 100 or 200 pg/ml DOX for 48 hours. 2×10^5^ BaF3 cells were then washed in PBS, resuspended in IL-3- free medium supplemented with the same concentration of DOX and 0.06 U/ml human erythropoietin and counted as described above.

### Immunoprecipitation and Immunoblotting

To assess protein phosphorylation, BaF3 cells and their derivatives were first starved in RPMI-10 IL-3-free media for 3 hrs at 37°C and were then acutely stimulated with 1 U/mL EPO for 10 min at 37°C. Cells were then washed twice with ice-cold PBS containing 1 mM phenylmethylsulfonyl fluoride (PMSF). For phosphotyrosine and phospho-protein blots, 1X HALT Protease and Phosphatase Inhibitor Cocktail (Thermo Scientific) and 500 µM hydrogen peroxide-activated sodium metavanadate were also added to the wash solution. Cells were lysed in FLAG-lysis buffer (50 mM Tris pH 7.4, 150 mM NaCl, 1 mM EDTA, 1% Triton-100) supplemented with protease and phosphatase inhibitors as described above. All lysates were incubated on ice for 20 minutes, followed by centrifugation at 14,000 rpm for 30 minutes at 4°C. The total protein concentration of the supernatants was determined using a bicinchoninic acid (BCA) protein assay kit (Pierce).

To immunoprecipitate FLAG-tagged CHAMP peptides, 50 µl of anti-FLAG M2 matrix gel (Sigma-Aldrich) was added to 0.5 mg of total protein and rotated overnight at 4°C. Immunoprecipitated samples were washed four times with 1 mL NET-N buffer (100 mM NaCl, 0.1 mM EDTA, 20 mM Tris-HCl pH 8.0, 0.1% Nonidet P-40) supplemented with protease inhibitors as above, pelleted and re-suspended in 2x Laemmli sample buffer (2x SB) supplemented with 200 mM dithiothreitol (DTT) and 5% β-mercaptoethanol (β-ME). Precipitated proteins and whole cell lysates were heated at 95°C for 5 min and then resolved by SDS-PAGE on either 7.5%, 10%, or 20% polyacrylamide gels according to the size of the protein of interest. The resolving gel was then transferred by electrophoresis to a 0.2 µm nitrocellulose or PVDF membrane. 0.09% SDS was added to the transfer buffer for membranes used to detect phosphorylated proteins.

Membranes were blocked with gentle agitation for two hours at room temperature in 5% nonfat dry milk/TBST. To detect the phosphorylated forms of JAK2 and STAT5, anti-phospho- JAK2 (Tyr1008) (clone D4A8, Cell Signaling) and anti-phospho-STAT5 (Y694) #9351 (Cell Signaling) were used. To detect the total JAK2 and STAT5, anti-JAK2 (clone D2E12, Cell Signaling) and anti-STAT5 #9363 (Cell Signaling) were used. An HRP-conjugated mouse anti-HA (clone 6E2, Cell Signaling) was used to detect the HA-tagged EPOR and all EPOR mutants. All antibodies were used at 1:1000 dilution. Membranes were incubated overnight with gentle agitation in primary antibody at 4°C, washed five times in TBST, and then incubated with gentle agitation for one hour at room temperature in a 1:10,000 dilution of donkey anti-mouse or donkey anti-rabbit HRP (Jackson Immunoresearch), as appropriate. To re-probe membranes, they were stripped in Restore Western Stripping Buffer (Thermo Scientific) for 15 min at room temperature with gentle agitation, washed five times in TBST, blocked in 5% milk/TBST for one hour at room temperature, and incubated overnight at 4°C with antibody, as described above. Membranes were incubated with Super Signal West Pico or Femto Chemiluminescent Substrates (Pierce) to detect protein bands.

### Split GFP Complementation Assay

BaF3 cells were transduced to express GFP1-10 fragment fused to EpoR, GFP11 fragment fused to CHAMP-1, or both. The EpoR-GFP1-10 fusion protein consists of (from N-terminus to C-terminus) the residue 1-258 from hEpoR or residue 1-257 from mEpoR, a 10-amino-acid flexible linker (GGSGGGGSGG), and the GFP1-10 segment. The GFP11-N1 fusion proteins consists of (from N-terminus to C-terminus) a FLAG tag, the GFP11 segment (residue 216-231), a GGG linker, and the CHAMP-1 sequence. For flow cytometry, 5×10^5^ cells were collected by centrifugation at 1,000 rpm for 10 minutes at 4°C, washed in cold PBS, and re-suspended in 300ul cold PBS. Cells were then analyzed by Cytoflex using green laser and plotted in FlowJo.

## Data availability

Chemical shift data has been uploaded to BMRB, with entry assigned accession number 51401.

## Acknowledgements

MM was supported by the HHMI Gilliam Fellowship. We thank Ross Federman and Lisa Petti for constructs. This work was supported by grants from the NIH to W.F.D. (R35-122603) and D.D. (CA037157). We thank Dr. Mark Kelly for technical assistance at the UCSF NMR Lab. This work used the QB3-Berkeley 900 MHz NMR Facility, which is supported by NIH GM68933.

## Author Contributions

M.M. and H.H. performed the computational design. L.H., W.B., and A.E. performed all experiments in cultured mouse cells. M.M., H.H., and H.J. synthesized and purified the peptides. M.M. and H.H. executed thiol-disulfide exchange experiments. M.M. and S.E.N. conducted the fluorescence quenching experiments. M.M., H.T.K., and Y.W. performed the NMR experiments. M.M., L.H., D.D., and W.F.D. designed the experiments, analyzed the data, and wrote the manuscript.

---

**Figure S1.**
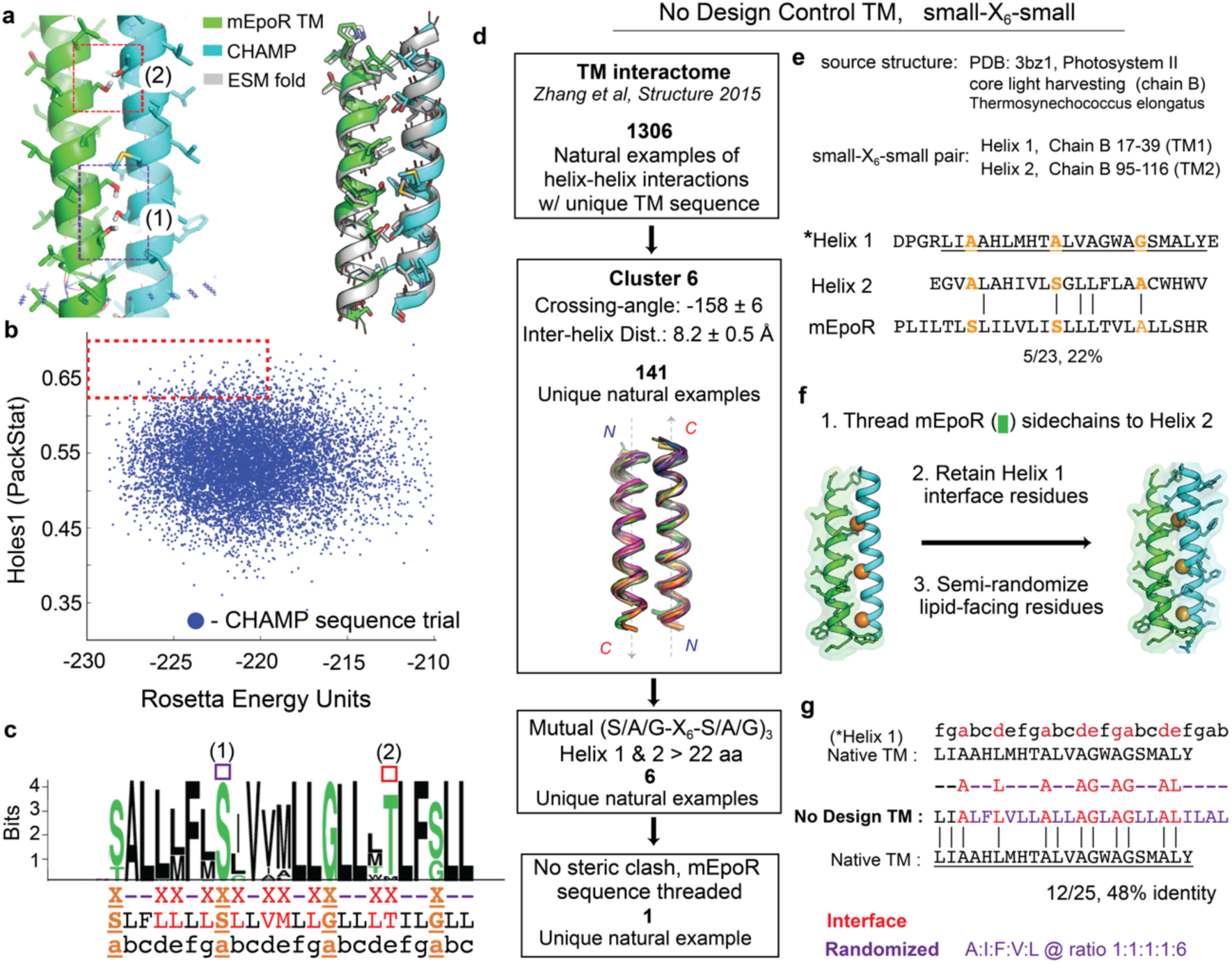
Protein design and ranking of output anti-mEpoR CHAMP sequences. (a) Left, final model of CHAMP-1/mEpoR TM domain complex of anti-parallel helices (green, mEpoR; cyan, CHAMP) from Rosetta showing two hydrogen inter-helical bond networks. Right, ESMfold *ab initio* predicted model of the complex (gray) overlaid with the Rosetta model, showing consistent prediction of register and sidechain interactions. The ESMfold model was predicted using a long glycine linker between the TM domains: (b) Scatter plot of each sequence design trial’s model Rosetta energy and packing voids score (RosettaHoles1^44^ via the “packstat” filter in RosettaScripts^43^). Only sequences with Rosetta energy greater than the mean energy of all designs were considered, and of those, only those having Holes scores in the top 10% were considered (Red dashed box). These top unique designed sequences were clustered based on sequence identity of interfacial residues, resulting in two unique sequences: the final CHAMP-1 and CHAMP-1 designs. (c) Sequence Logo of all sequence output from CHAMP design. Below shows design protocol, with residues that were fixed or allowed to sample identities/rotamers: black hyphen, fixed to a semi-random apolar amion acid; red X, any lipid-friendly residue (GATSVLIFM); orange X, designated small-X_6_-small positions (GAS). (d) The “No Design” control TM sequence was filtered from all possible helical pairs from out previous database to just the anti-parallel left-handed close-packing motif structural motif (small-X_6_-small consensus sequence) with sequences having mutual small-X_6_-small motifs repeated three times, as in mEpoR and CHAMP-1, and being at least 22 residue per helix. Six helix pairs matched these criteria. For each helix within the helix pair molecular model, the mEpoR sequence was threaded onto one helix in register with the small-X_6_-small motif and mEpoR’s sidechain steric overlap with the alternate native sequence (fixed rotamers) was assessed. Only in 1 of the 12 cases was there no steric clashes. (e) Source structure and TM domain antiparallel helix pair sequence of “No Design” control TM, Photosystem II light harvesting complex (PDB:3bz1, chain B), comprised of TM 1 and 2. Helix 2, on which mEpoR was threaded, only has 22% sequence identity with mEpoR, which essentially random given the statistical distribution of amino acids in membrane proteins. (f) Sequence selection for the “No Design” control TM. A suitable helix pair was extracted from a natural protein. Residue identities at the helix-helix interface were retained, including the small-X_6_-small motif (orange spheres). Lipid-facing residues were altered. (g) Final sequence selection and comparison of the “No Design” control TM to the Photosystem II source TM domain

**Figure S2.**
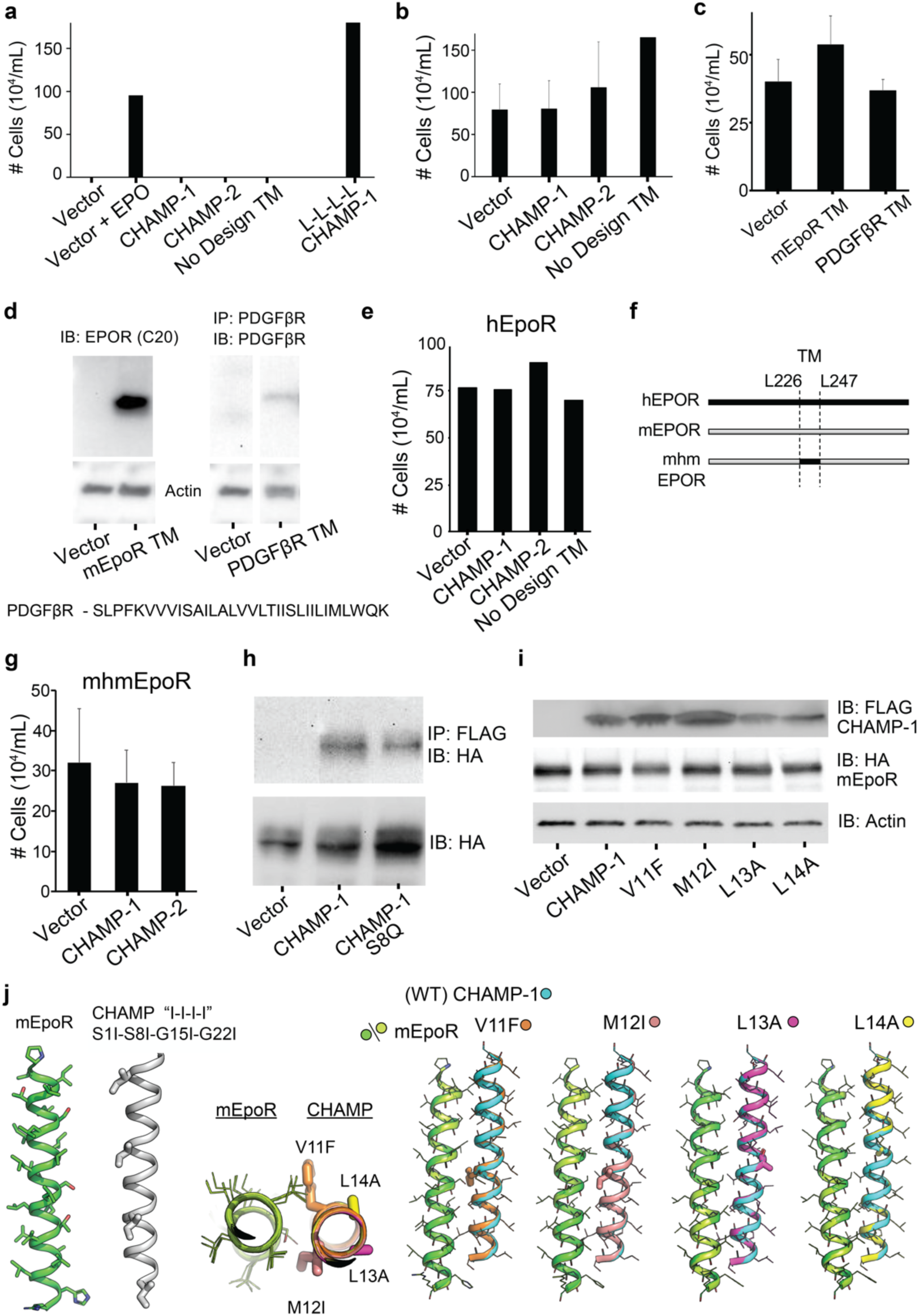
Effects of expressed exogenous TM domain on EpoR expressing BaF3 cells. (a) BaF3/mEpoR cells transduced with empty vector or a TM proteins (CHAMP-1, CHAMP-1, No Design TM, or CHAMP-1 mutant “L-L-L-L”) grown in media not treated with IL-3 and EPO. The number of live cells at day 4 is shown. EPO-treated cells transduced only empty vector are the positive control for EPOR-dependent proliferation. CHAMP-1 “L-L-L-L” mutant causes EPO-independent proliferation, but the other TM proteins do not. (b) BaF3/mEpoR cell counts on day 4 upon stably expressing empty vector, CHAMP-1, CHAMP-1, or No Design TM and supplementing with 5% WEHI-conditioned medium as source of IL-3. (average and standard error as error bars of three replicate experiments except for No Design TM, which was tested once). (c) BaF3/mEpoR cells expressing empty vector, a short mEpoR TM domain protein construct, or a previous published PDGFβR TM domain construct (Ref 46) were incubated for eight days in medium supplemented with 0.06 U/mL EPO. The average number of live cells is shown with standard error of three independent experiments. (d) Upper, western blots of extracts of BaF3/mEpoR cells stably expressing the mEpoR TM construct (probed with antibody targeting the C20 cytoplasmic epitope) and PDGFβR TM construct. The PDGFβR TM construct was first immunoprecipitated to increase concentration due to low expression level then immunoblotted, using rabbit anti- serum targeting a PDGFβR C-terminal cytoplasmic region epitope as in Ref 46. Bottom, PDGFβR receptor TM domain sequence. Full sequences of expressed protein constructs are listed in Table S1. (e) BaF3/hEPOR cells expressing empty vector, CHAMP-1, CHAMP-1, or No Design TM were incubated for four days in medium supplemented with 0.06 U/mL EPO. The number of live cells is shown. (f) mhmEpoR construct having the cytoplasmic and extracellular domain sequences of mEpoR but the TM domain of hEpoR (sequence in Table S1). (g) BaF3/mhmEpoR cells expressing empty vector, CHAMP-1, CHAMP-1, or No Design TM were incubated for eight days in medium supplemented with 0.06 U/mL EPO. The average number of live cells is shown with standard error of three independent experiments. (h) Extracts from BaF3/mEpoR expressing empty vector or FLAG-tagged CHAMP-1 or mutant SQ8 were immunoprecipitated with anti-FLAG, subjected to SDS-PAGE, and probed with anti-HA to detect HA-tagged mEpoR. Bottom panel shows samples without prior immunoprecipitation. (i) Western blots of extracts of BaF3/mEpoR cells stably expressing wildtype or mutant CHAMP-1. Blots were probed with antibodies recognizing CHAMP (anti-FLAG), mEpoR (anti-HA), and actin as a loading control. (j) ESMfold predictions of CHAMP-1 mutants with EpoR TM domain. Left, the prediction of mEpoR (green) with CHAMP-1 “I-I-I-I” mutant (gray, Ile in sticks) results in two distant non-interacting helices. Right, overlay of ESMfold predictions for the EpoR/CHAMP-1 complex (green and blue helices, respectively) with predictions for the complex baring point mutations on CHAMP-1: V11F (orange), M12I (pink), L13A (magenta), L14A (yellow). Predictions show the lowest energy structure is similar to the initial design model for each point mutant.

**Figure S3.**
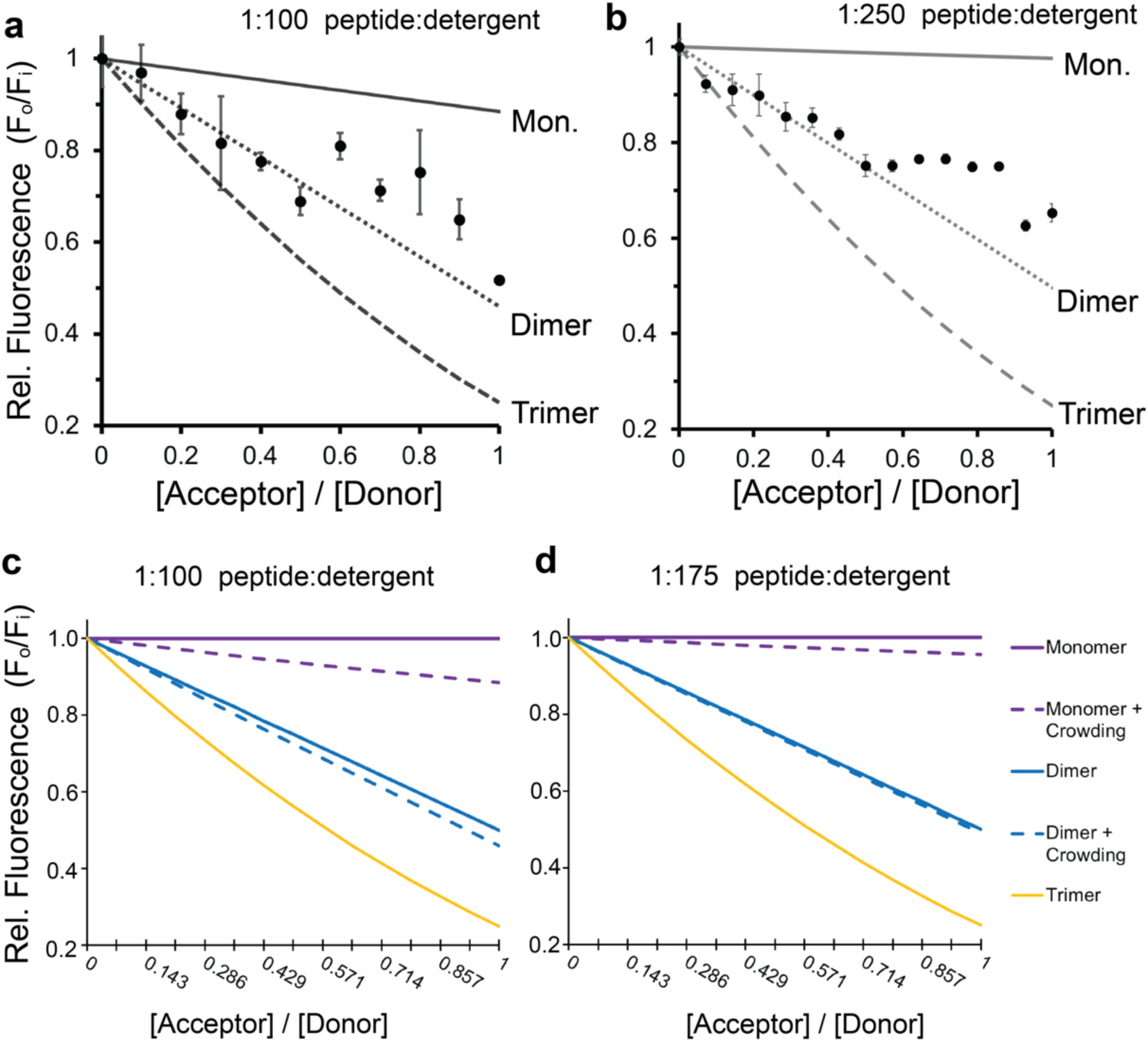
Fluorescence quenching of labeled-mEpoR by CHAMP-1 in C14B micelles at different peptide:detergent ratios and theoretical FRET from non-specific peptide crowding effects. Relative (Rel.) 460 nm fluorescence emission of 1.5 µM 7-diethylamino-4-methylcoumerin-labeled mEpoR TM peptide shows FRET quenching as fluorescein-labeled CHAMP-1 is titrated in C14-betaine, 50 mM Tris-HCl pH 8, 100 mM NaCl, 0.5 mM EDTA, 5 mM TCEP buffer at fixed equimolar total peptide concentration (n=3-6; bars, standard error) (a) using a detergent to total peptide ratio of 100:1 (1.1 micelles per total peptide) or (b) Using a detergent to total peptide ratio of 250:1 (2.8 micelles per total peptide). Theoretical FRET curves expected for monomeric (Mon.), dimeric, and trimeric complexes are plotted, which include a micelle crowding factor (gray) to account for the non-specific contribution to FRET quenching from co-habitation of donor and quenching peptides in the same micelle at random, dependent upon peptide to detergent relative concentrations (mole fractions). The vendor’s micelle aggregation number of 83 detergent molecules per micelle and a Poisson distribution were used to calculate the expected population of dimer to hexamer given the relative moles of peptides and micelles in solution. Each population fraction was used to compute the expected non-specific FRET quenching contribution due to each species dimer to hexamer, which can be added to the model of FRET quenching expected from specific donor:acceptor molecule interactions for a given stoichiometry or species. Theoretical model of FRET quenching for monomer, dimer, and trimeric association with the contribution of non-specific FRET from higher-order oligomers (dashed line) and without the crowding factor (solid) at (c) 1:100 peptide to C14-B mole ratio and (d) 1:175 peptide to C14B- mol ratio.

**Figure S4.**
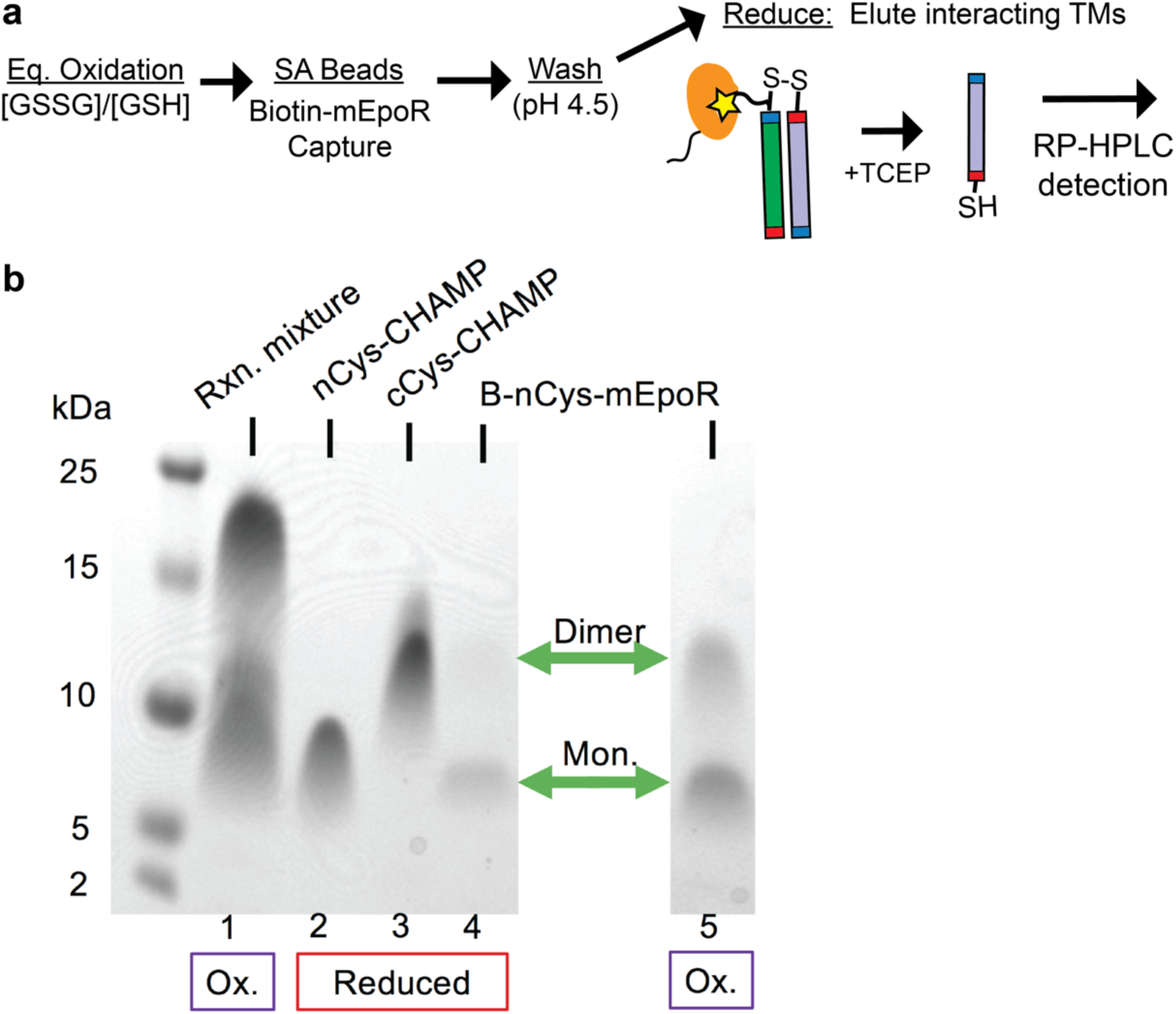
Schematic and SDS-PAGE of the three peptide thiol-disulfide exchange with biotin capture. **(a)** Workflow for equilibrium thiol-disulfide exchange modified for biotin capture and isolation of only mEpoR- containing disulfide bonded TM dimers, followed by reduction and quantification of the interacting TM helices by reverse phase high-performance liquid chromatography (RP-HPLC). SA, Streptavidin; GSSG, GSH, oxidized and reduced glutathione, respectively; Eq., equilibrium. **(b)** Sodium dodecyl sulfate polyacrylamide gel electrophoresis (SDS-PAGE) of equilibrated oxidized TM peptides in 20 mM m*yristyl sulfobetaine* (C14B) micelles (lanes 1, 5), or purified peptides (lanes 2, 3, 4) reduced in 10 molar equivalents of tris(2-carboxyethyl)phosphine (TCEP). *Lane 1*, mixture of 100 µM Biotinylated nCys-mEpoR with 4-fold molar excess of CHAMP – 2 equivalents of each nCys- CHAMP1 and cCys-CHAMP1 V2 oxidized overnight at a molar ratio of oxidized to reduced glutathione of 0.2. *Lane* 5, 100 µM Biotinylated nCys-mEpoR alone oxidized under the same conditions. Green arrows denote mEpoR monomer and dimer bands. Ox., oxidizing conditions; Mon., Monomer; kDa, kilodaltons; Rxn., Reaction.

**Figure S5.**
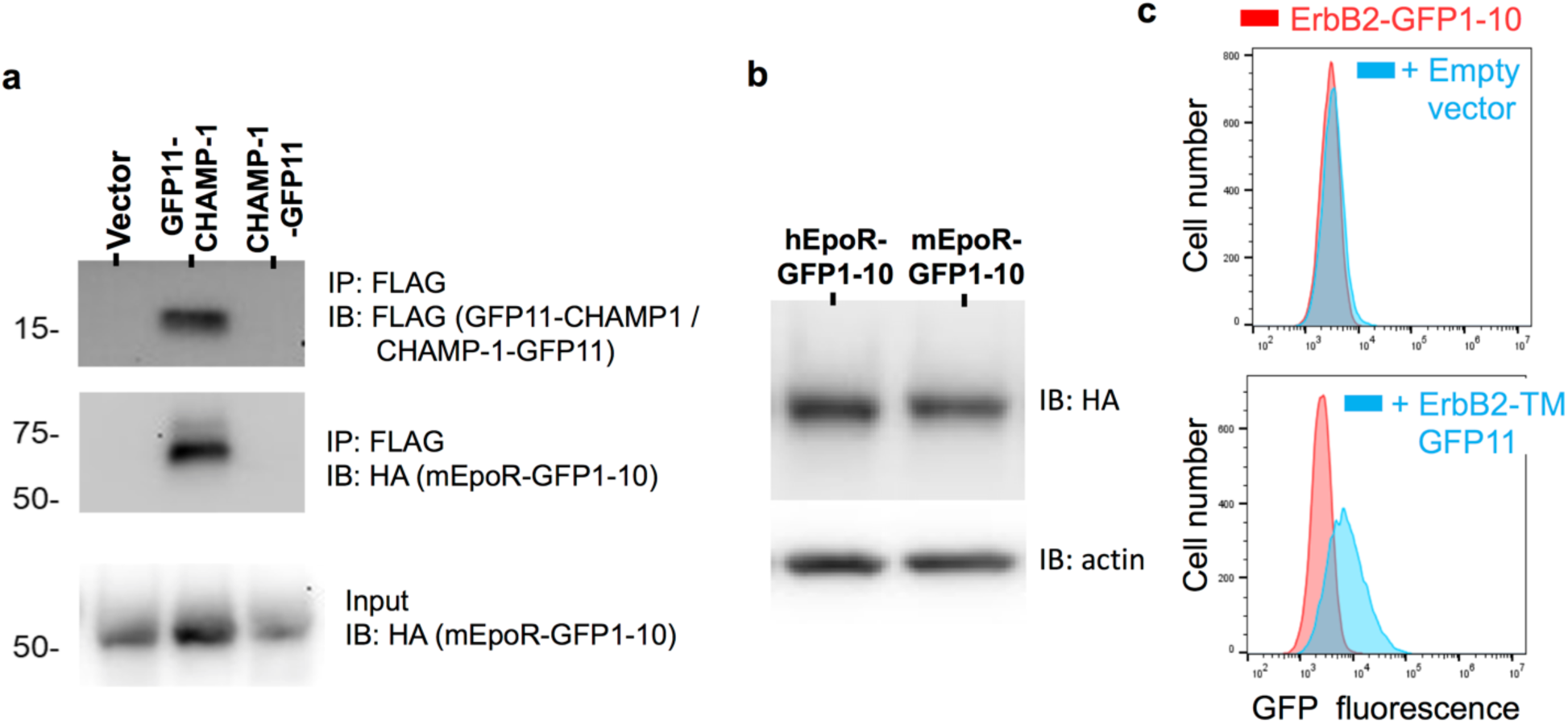
Control split GFP protein complement experiments. **(a)** Extracts were prepared from mEpoR-GFP1-10 cells expressing empty vector or FLAG-tagged CHAMP-1 with a fused N-terminal GFP11 (GFP11-CHAMP-1) or a fused C-terminal GFP11 (CHAMP-1-GFP11). Extracts were subjected to SDS-PAGE directly (input) or after immunoprecipitation with anti-FLAG antibody, which recognizes both FLAG-tagged CHAMP-1 proteins. Blots were probed with antibodies that recognize FLAG to detect the CHAMP-1 proteins or HA to detect HA-tagged mEpoR-GFP1-10. **(b)** Extracts from cells expressing either the mEpoR or the hEpoR C-terminally fused to GFP1-10 were subjected to SDS-PAGE and immunoblotted with antibodies that recognize HA fused to EpoR or actin as a loading control. The numbers below the lanes are the relative expression of the EpoR normalized to actin. **(c)** BaF3 (tTa) cells expressing ErbB2 C-terminally fused to GFP1-10 transduced with empty pTight vector or pTight containing the TM domain of ErbB2 fused to GFP11 (ErbB2TM-GFP11). Flow cytometry was performed on cells incubated in 1 nM doxycycline to repress ErbB2TM-GFP11 (top panel) or in 1 pM doxycycline (bottom panel).

**Figure S6.**
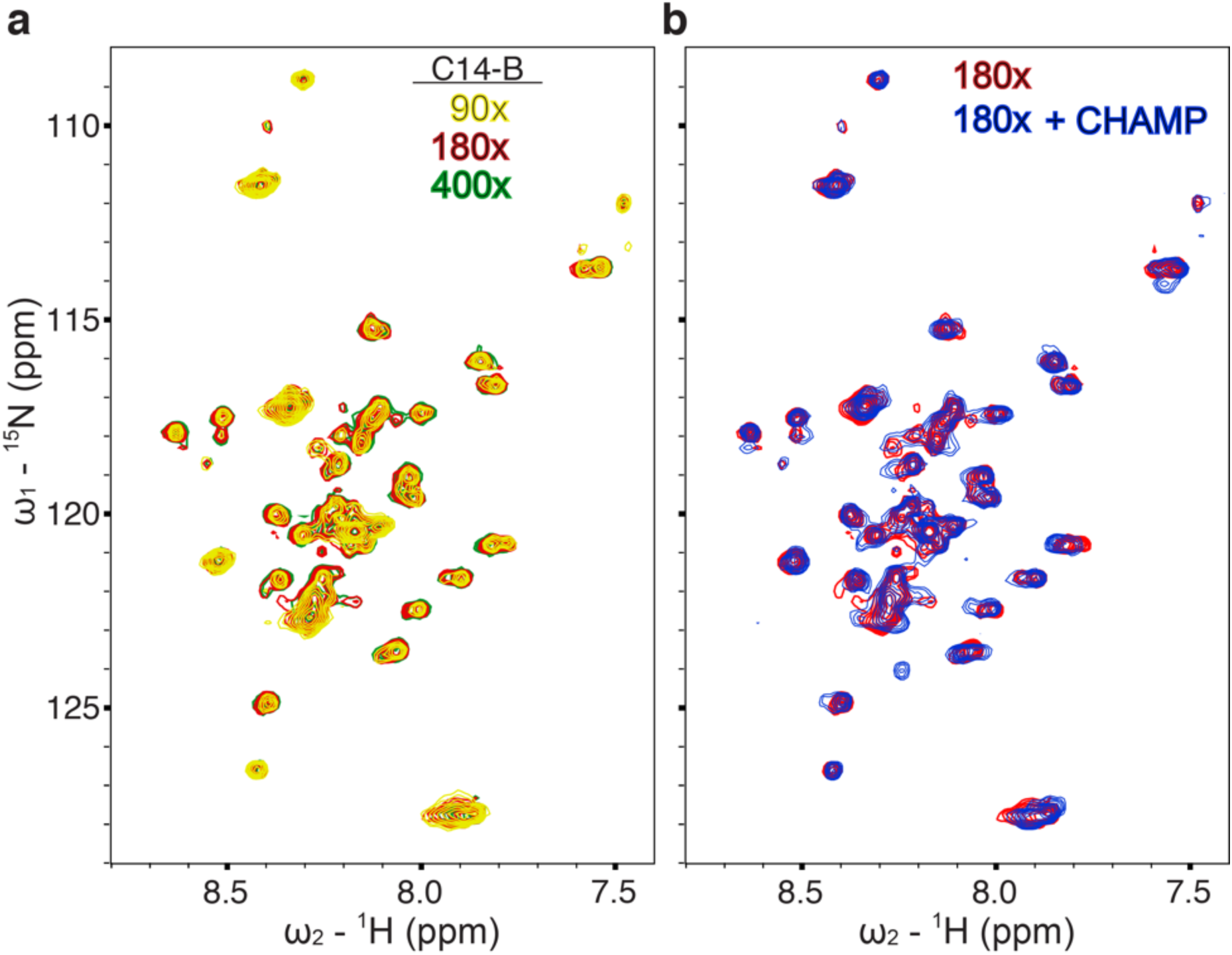
1^5^N mEpoR-TM1 monomer-homodimer behavior in C14-Betaine micelles and binding to CHAMP- 1. (a) ^1^H-^15^N HSQC spectra of 0.3 µM ^15^N mEpoR-TM1 when diluted in myristyl-sulfobetaine (C14-B) micelles at 90, 180, and 400 molar equivalents of detergent to peptide (27 mM, yellow; 54 mM, red; 120 mM, blue) in 40 mM sodium acetate buffer pH 5.2, 20 mM NaCl, 0.5 mM EDTA, 5 mM DTT at 45° C and 800 MHz. C14-B ∼90 detergent molecules per micelle. Multiple populations are observed which show resonance-dependent peak broadening over the titration, but negligible chemical shift perturbations. **(b)** Maintaining both the same ^15^N mEpoR-TM1 concentration (0.3 µM) and the total detergent:protein molar ratio (180), 1.35 molar equivalents of unlabeled CHAMP-1 were added and the ^1^H-^15^N HSQC spectra is shown overlaid (Blue, 0.4 µM CHAMP, 108 mM C14-B; 360x C14-B per mole mEpoR-TM1). Many new peaks are observed and some existing peaks show large decays in intensity, indicative of mEpoR-TM1’s interaction with CHAMP-1 slow on the NMR timescale.

**Figure S7.**
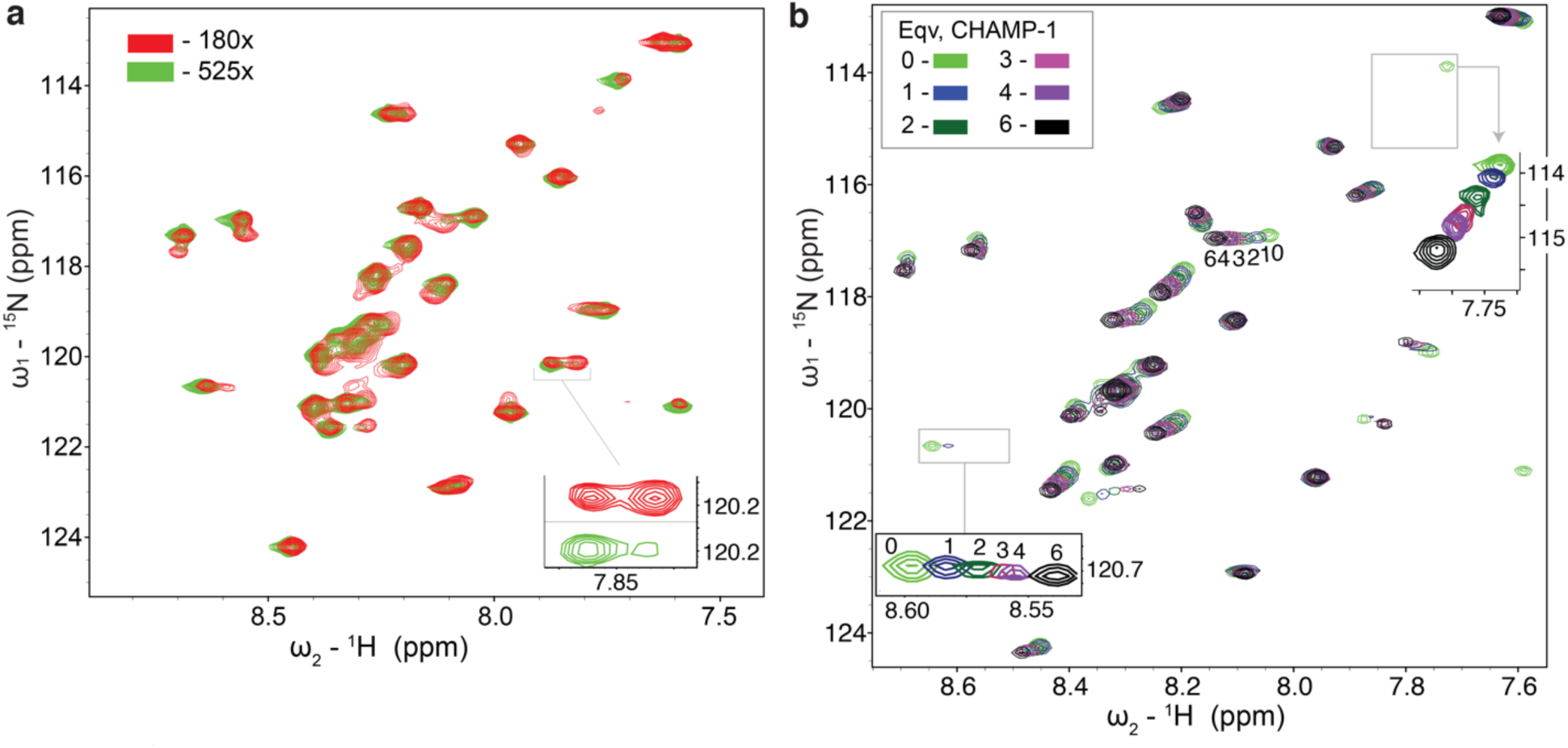
^15^N mEpoR-TM2 monomer-homodimer behavior in C14-Betaine micelles and binding to CHAMP- 1. (a) The ^1^H-^15^N HSQC spectra of 0.3 µM ^15^N mEpoR-TM2 at 45° C, pH 5.2, and 800 MHz when diluted in myristyl-sulfobetaine (C14-B) micelles at 180 and 520 molar equivalents of detergent (54 mM, red; 156 mM, green). The more concentrated sample (180x, red) shows a strong second set of peaks emerging in exchange that is slow on the NMR time scale, likely representing the mEpoR-TM2 in the homodimeric state. *Right inset*, example pair of related, interconverting peaks dependent on the detergent concentration. (b) Titration of 1, 2, 3, 4, and 6 molar equivalents (eqv.) of CHAMP (navy, dark green, red, purple, black, respectively) to 0.3 µM ^15^N mEpoR-TM2 where additional C14-B is added alongside each eqv. CHAMP to maintain a constant molar ratio of C14-B to total peptide of 180:1. Reference spectra (no CHAMP, green) in (b) is predominantly the monomeric species at 520:1 ratio C14-B:mEpoR- TM2. Insets track shifting resonances at lower contour.

**Figure S8.**
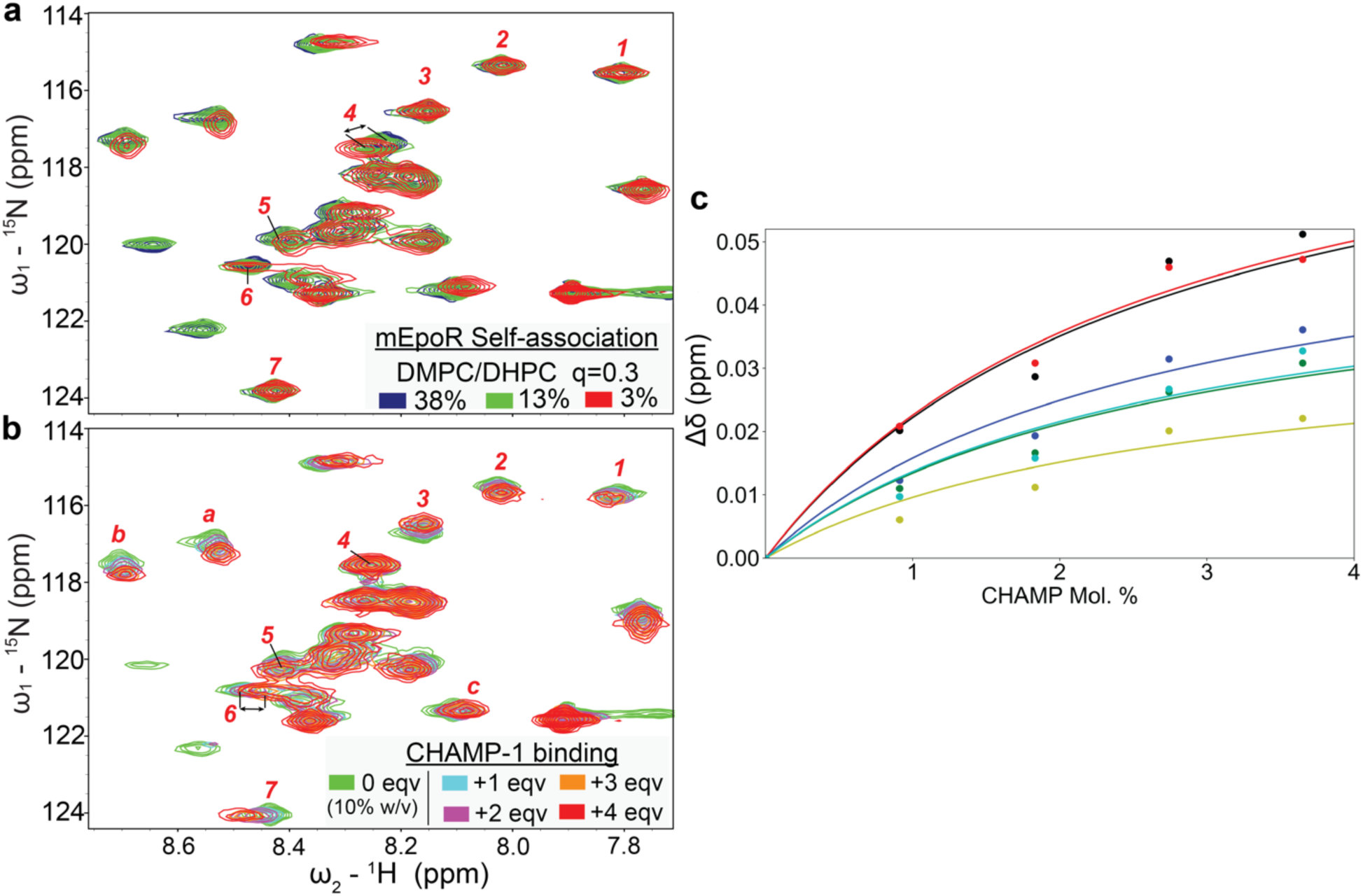
mEpoR-TM2 monomer-homodimer equilibrium and binding to CHAMP-1 in DHPC/DMPC q=0.3 bicelles. ^1^H-^15^N HQSC spectral changes from 0.5 mM ^15^N-mEpoR-TM2 at 45° C, pH 5.2, and 800 MHz upon (**a**) titration of DMPC/DMPC q=0.3 bicelles at 38%, 13%, and 3% (w/v) at constant mEpoR-TM2 (navy, green, red, respectively) or (**b**) titration of 1, 2, 3, and 4 molar equivalent (eqv.) unlabeled CHAMP (cyan, orange, maroon, red, respectively) at a constant bicelle concentration of 10%, overlaid on a 10% bicelle (no CHAMP, green) reference spectrum. Both (**a**) the mEpoR homo-oligomer (likely homodimer) and (**b**) mEpoR-CHAMP heterodimer complex exhibit chemical shift perturbation relative to monomer resonances indicating a ‘fast’ chemical exchange. The numbered resonances denote induced chemical shift changes that differ significantly between the two titrations, suggesting two distinct states (**a**, mEpoR homodimer; **b** mEpoR-CHAMP heterodimer). Peaks 1, 2, 3, 6 and 7 do not shift as mEpoR converts from monomer to homodimer, while these peaks shift significantly upon CHAMP binding; peak 4 shifts upon homodimerization but not upon CHAMP binding; peak 5 shifts in different directions. Lowercase letters denote resonances experiencing peak shift similar between **a** and **b**. (**c**) Global fit to induced chemical shifts of resonances *a*, *b*, *c*, *1*, *2*, *5* upon CHAMP titration to calculate a binding affinity (K_d_) of 2.7 ± 0.6 mol %, in mol % peptide relative to DMPC.

**Figure S9.**
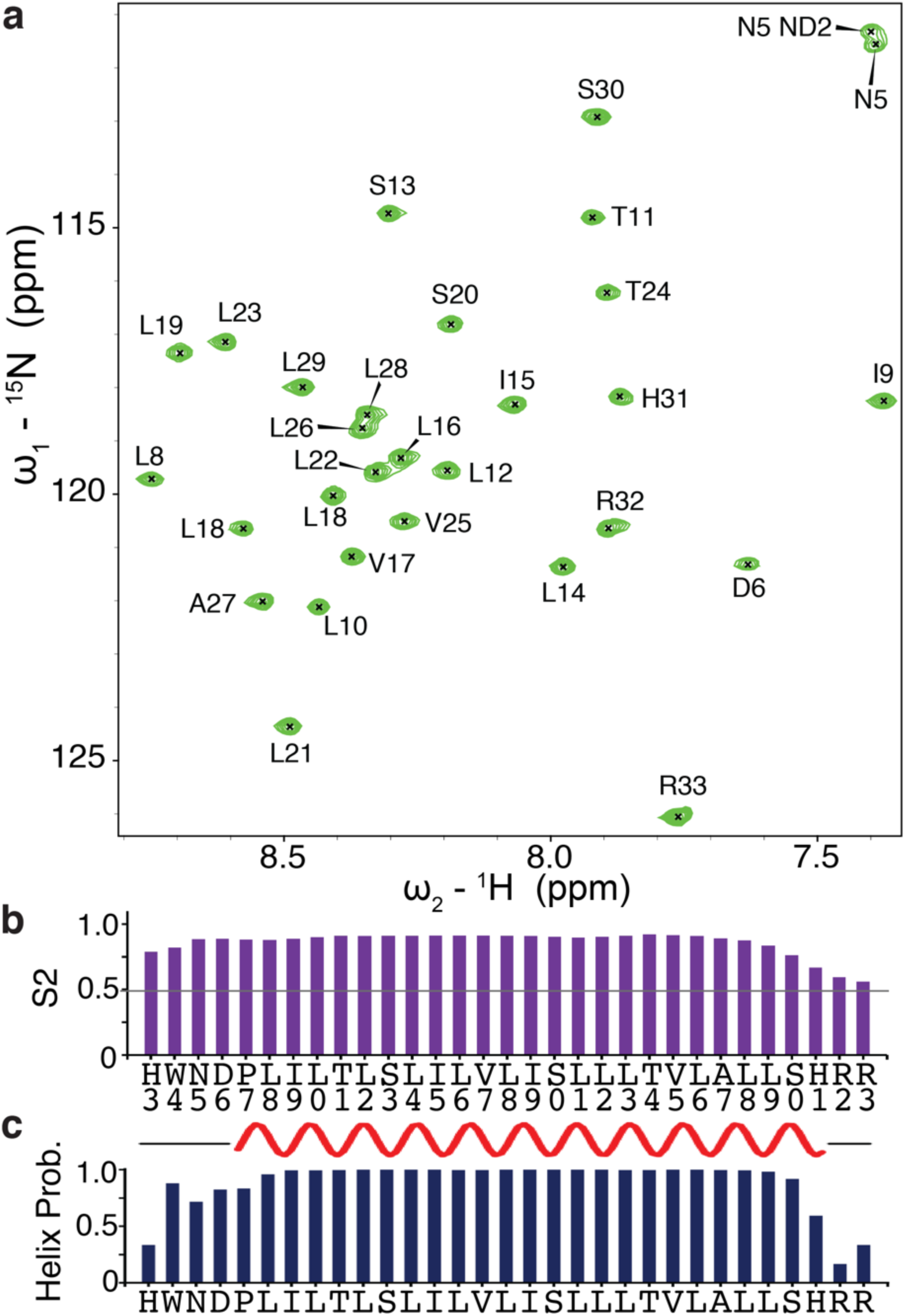
^1^H-^15^N HSQC spectra and per-residue structural features of monomeric mEpoR-TM2 in excess DPC. **(a)** Assigned ^1^H-^15^N HSQC spectra of 200 uM ^15^N mEpoR at 45° C, 800 MHz, pH 5.2, and high [DPC] : [protein] molar ratio: 600 or 12 micelles per protein. **(b)** predicted order parameter (S2) from TALOS+. **(c)** Predicted helicity from chemical shift index software. *Top*, predicted helical (red) and disordered (black) residues from CSI-3. *Bottom*, probability of helical secondary structure predicted by TALOS+. Prob., probability.

**Figure S10.**
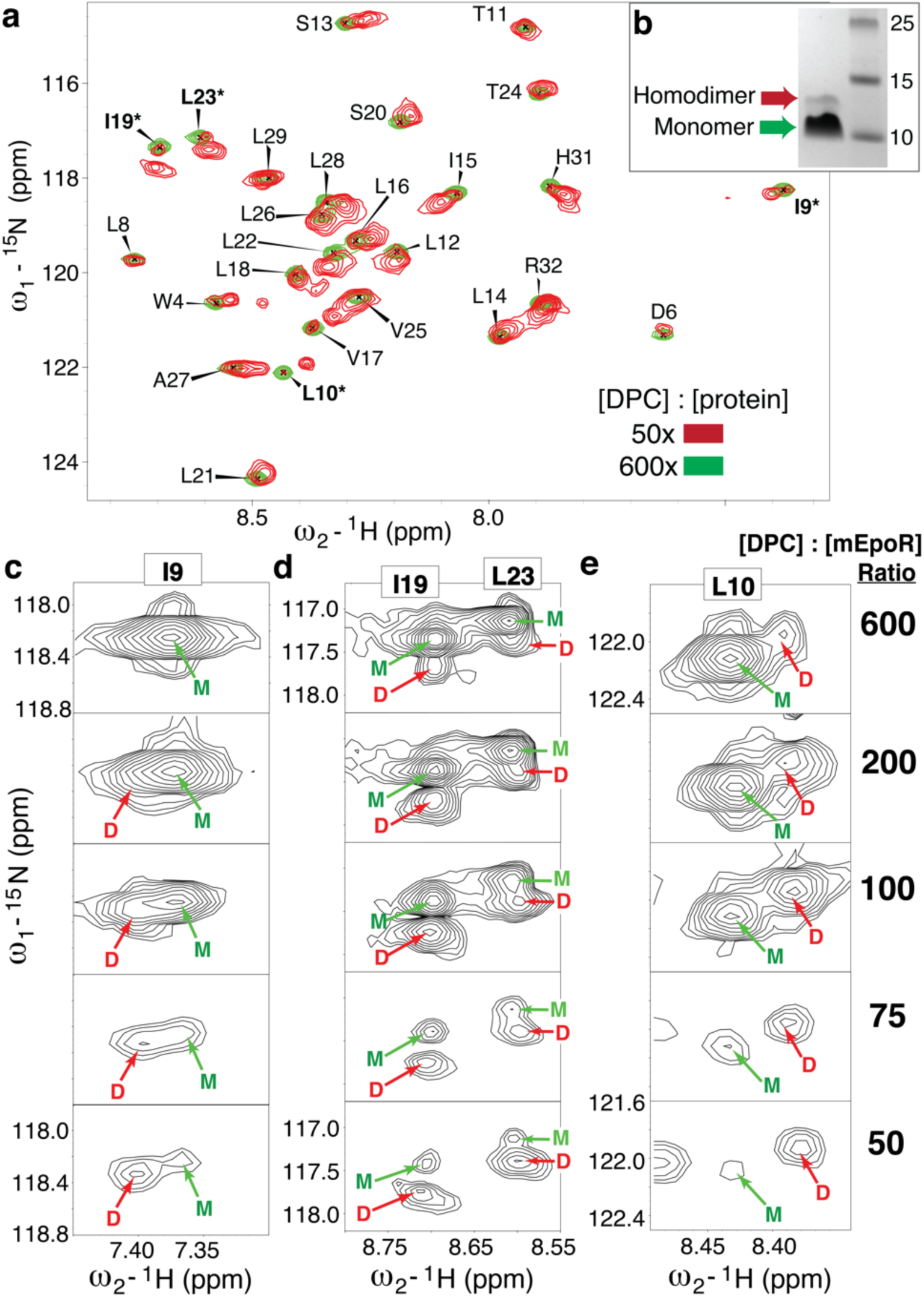
mEpoR Monomer-Homodimer equilibrium upon DPC titration (a). Assigned ^1^H-^15^N HSQC spectrum of 150 uM ^15^N mEpoR at high DPC molar ratio (600x, green) with the spectrum at low DPC concentration (50x, red) overlaid; the latter [DPC] concentration corresponds to approximately 1 TM peptide per 1 micelle. Recorded at 45° C, 800 MHz, pH 5.2, with the predominantly monomer spectra (green) contoured 10-fold higher than the largely homodimer spectra (red). **(b)** SDS-PAGE of mEpoR showing non-covalent monomer-dimer equilibrium. Mon, monomer. **(c-e)** Examples of newly emerging peaks (starred, bolded residues in **a**) of the mEpoR TM homodimer species upon reduction of DPC detergent concentration, titrated at 600, 200, 100, 75, and 50 molar ratios of detergent to TM peptide, showing the mEpoR TM peptide monomer-homodimer equilibria in the absence of CHAMP. Further concentrating mEpoR-TM2 to only 25x DPC molar ratio results in a sparse spectrum suggestive on peptide aggregation, where only 2 peaks (R32 and R33) are observed.

**Figure S11.**
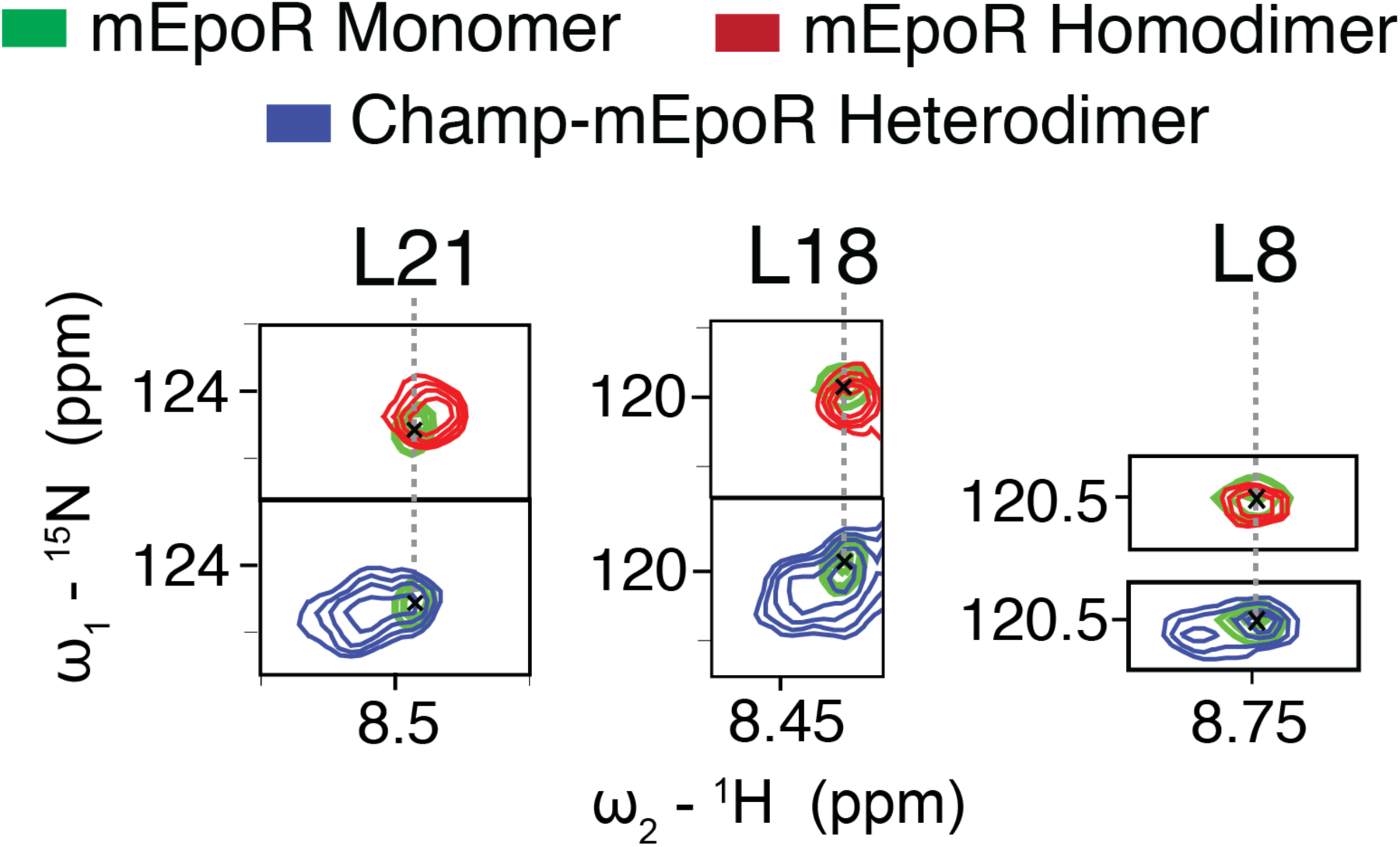
Characteristic ^1^H-^15^N HSQC peaks of mEpoR in the monomeric, homodimeric, and heterodimer (complex with CHAMP-1) states. ^1^H-^15^N HSQC spectrum of mEpoR-TM2 in its monomeric state (green, diluted in 600x mol excess DPC) is shown as a reference. Either increasing the mEpoR-TM2 concentration relative to detergent or adding CHAMP-1 peptide induce a second set of peaks corresponding to significant populations of slow-exchanging mEpoR dimeric species; full spectra are shown in Figures 5a and S10. Respectively, these correspond to the homodimeric assembly (red, more concentrated 50x mol excess DPC, i.e. about 1 micelle per mEpoR-TM2 peptide) and the heterodimeric complex with the CHAMP-1 peptide (blue, 7-fold molar excess CHAMP). Most of the ^1^H-^15^N induced chemical shift changes between mEpoR homodimer and heterodimer similar and nearly overlap. However, several notable resonances such as for Leu21 and Leu18 show distinct characteristic peak shifts (i.e. different directions) that distinguish the mEpoR homodimer and heterodimer state. All spectrum at 800 MHz, pH 5.2, and 45° C.

**Figure S12.**
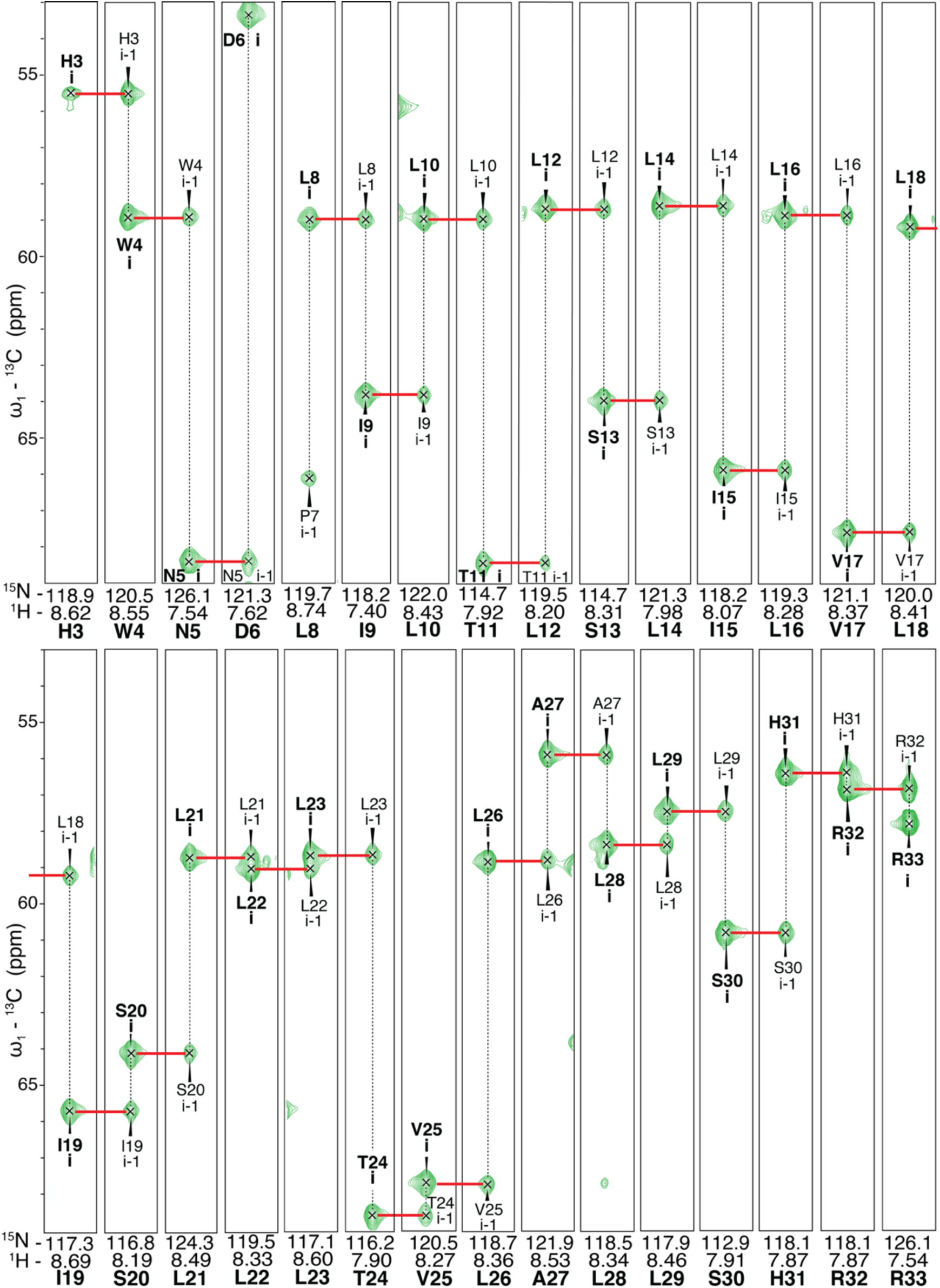
HNCA Assignment of mEpoR-TM2 monomer. ^1^H^13^C^15^N mEpoR at 1 mM in 400 mM ^2^H-DPC at 45° C, 800 MHz, pH 5.2. Red lines denote *i* and *i-1* resonance connectivity; X axis, ^15^N. Pro 7 assigned by L8 i-1 and lack of ^15^N-^1^H i peak or D6 i-1 peaks.

**Figure S13.**
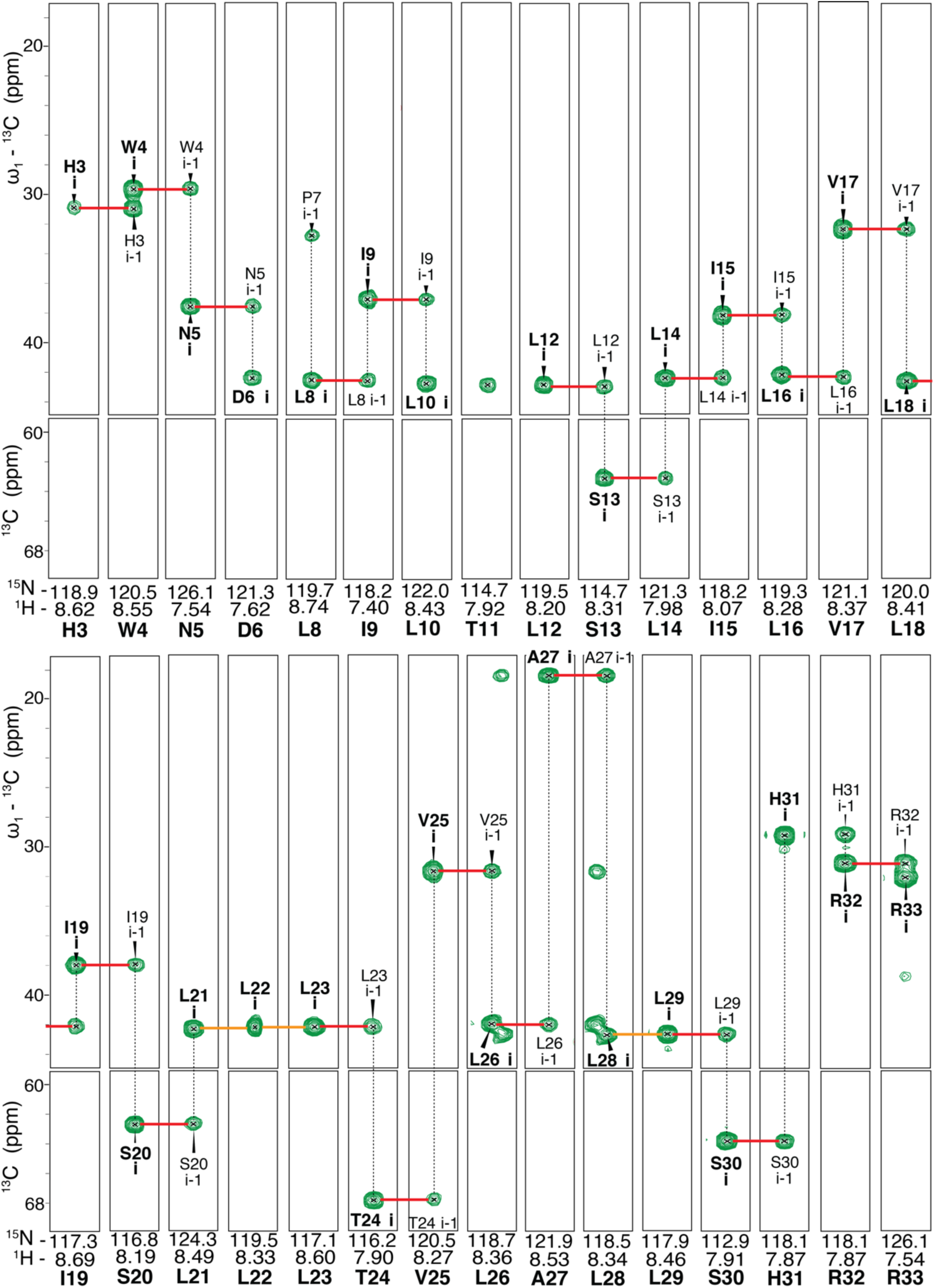
HNCB Assignment of mEpoR-TM2 monomer. ^1^H^13^C^15^N mEpoR at 1 mM in 400 mM ^2^H-DPC at 45° C, 800 MHz, pH 5.2. Orange lines denote ambiguous connectivity due to overlap of *i* and *i-1* resonances. Red lines denote *i* to *i-1* links. X axis, ^15^N. T11 displays no residue *i* CB-N resonance and L12 shows no *i-1* CB-N resonance to T11. Although a resonance exists in the T11 N-H strip, which is likely the L10 *i-1* CB-N resonance.

**Figure S14.**
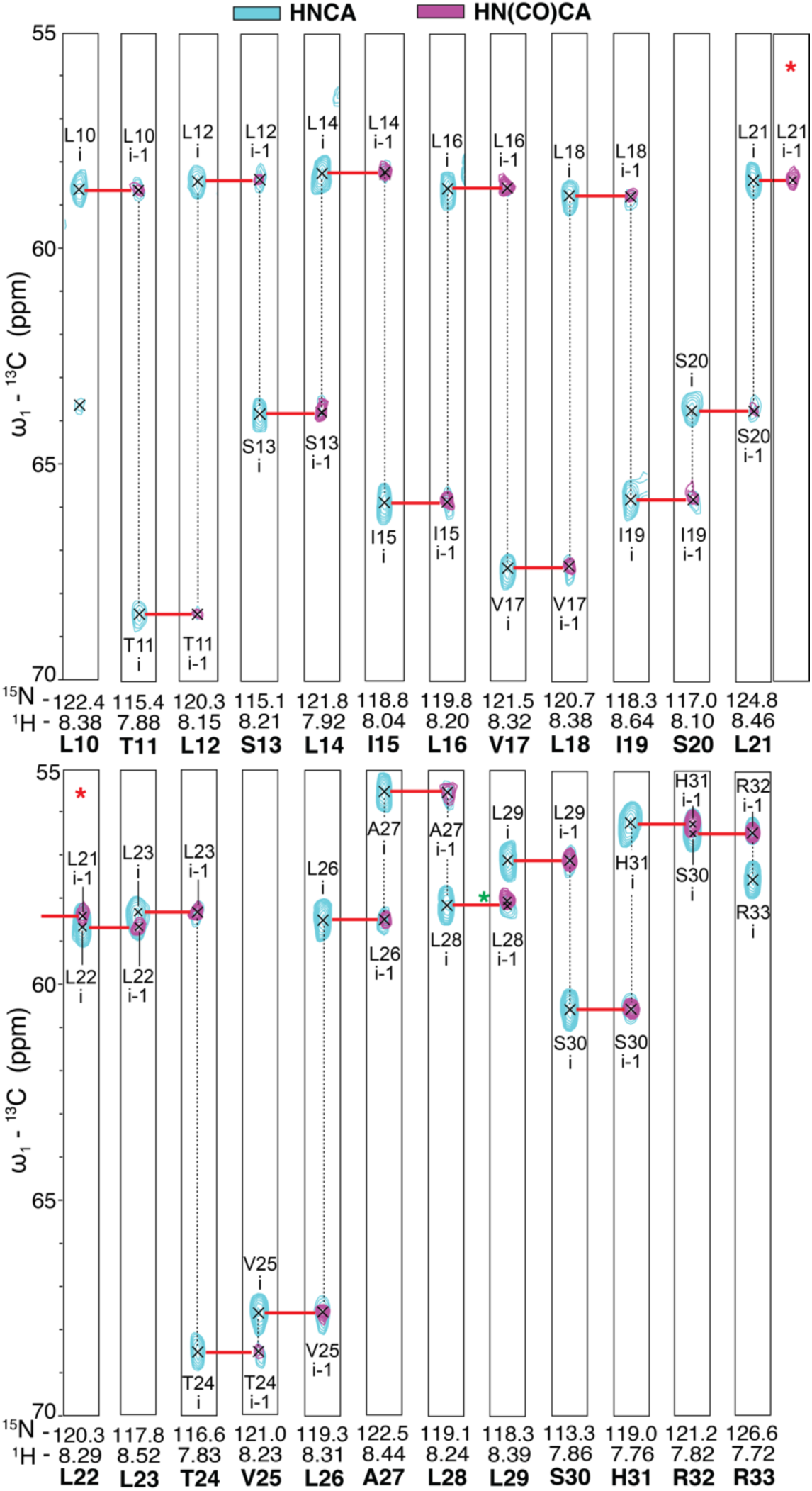
Paired HNCA and HNcoCA spectra of ^1^H^13^C^15^N mEpoR in complex with CHAMP-1. ^1^H^13^C^15^N mEpoR at 2 mM with 10 mM CHAMP-1 in 800 mM ^2^H-DPC at 45° C, 900 MHz, pH 5.2; HNCA, cyan; HNcoCA, magenta. Red lines denote *i* and *i-1* resonance connectivity; X axis, ^15^N. Red asterisk, repeated L22 strip of HCcoCA to show connectivity to L21 across rows of strips. Green asterisk, discrepancy in HNCA and HNcoCA peak position of L28 CA by 0.15 ppm (^13^C). H31, R32, and R33 resonances contoured 5, 10, 20 times higher, respectively.

**Figure S15.**
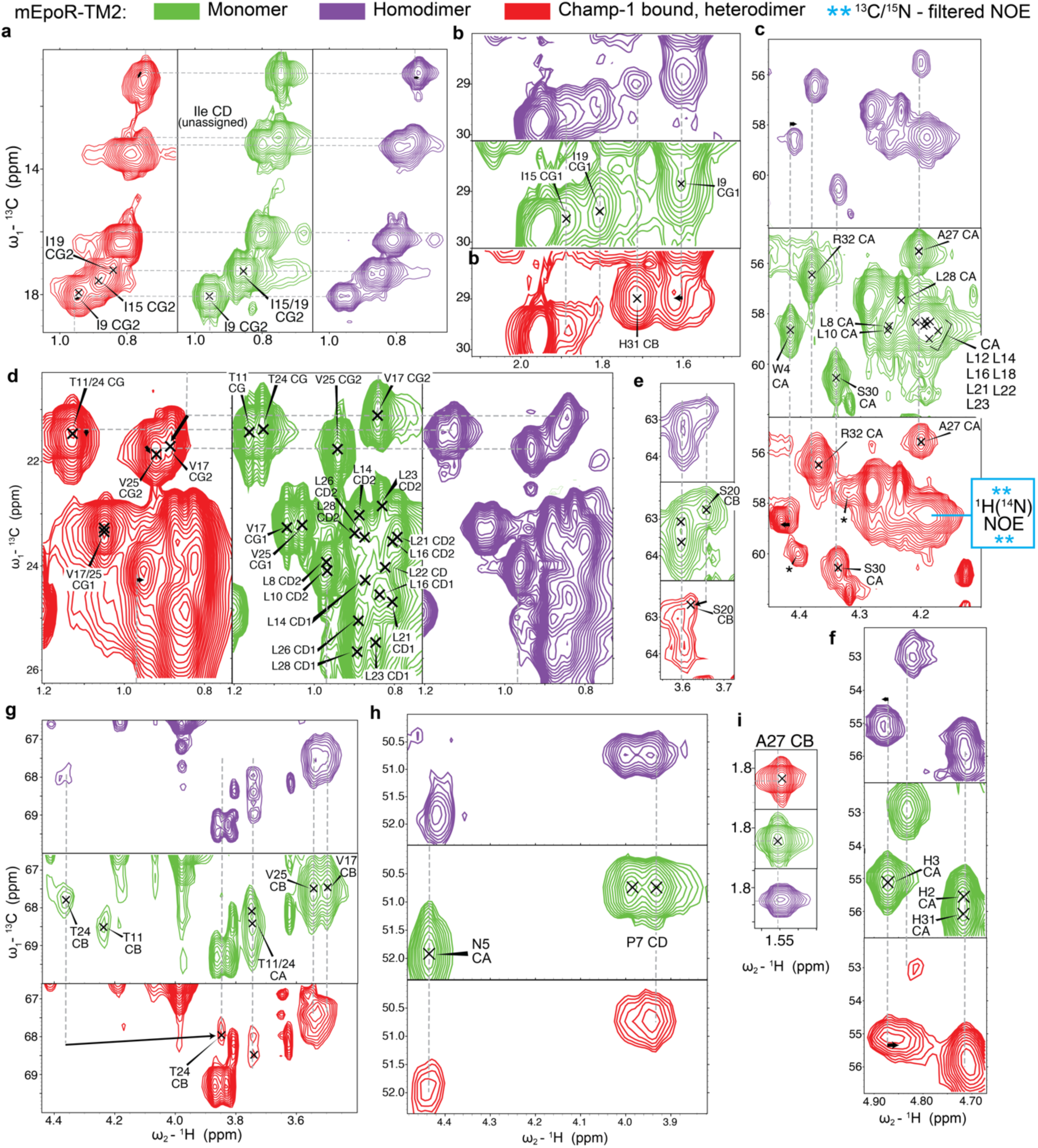
Comparison of ^1^H-^13^C HSQC spectra of mEpoR-TM2 from monomeric (green), CHAMP-1-bound (red), and partially homodimeric (purple) samples. (a-i) Different spectral regions perturbed. More monomeric resonances assigned from backbone-backbone and backbone sidechain experiments. Noted heterodimer assignments were derived independently by backbone-backbone, backbone-sidechain, and ^15^N-edited NOESY. *Cyan inset*, location of the isolated ^13^C resonance having a transfer NOE to a ^14^N-attached amide ^1^H peaks in the 13C-edited/^13^C,^15^N-filtered HSQC-NOESY (Fig. 5).

**Figure S16.**
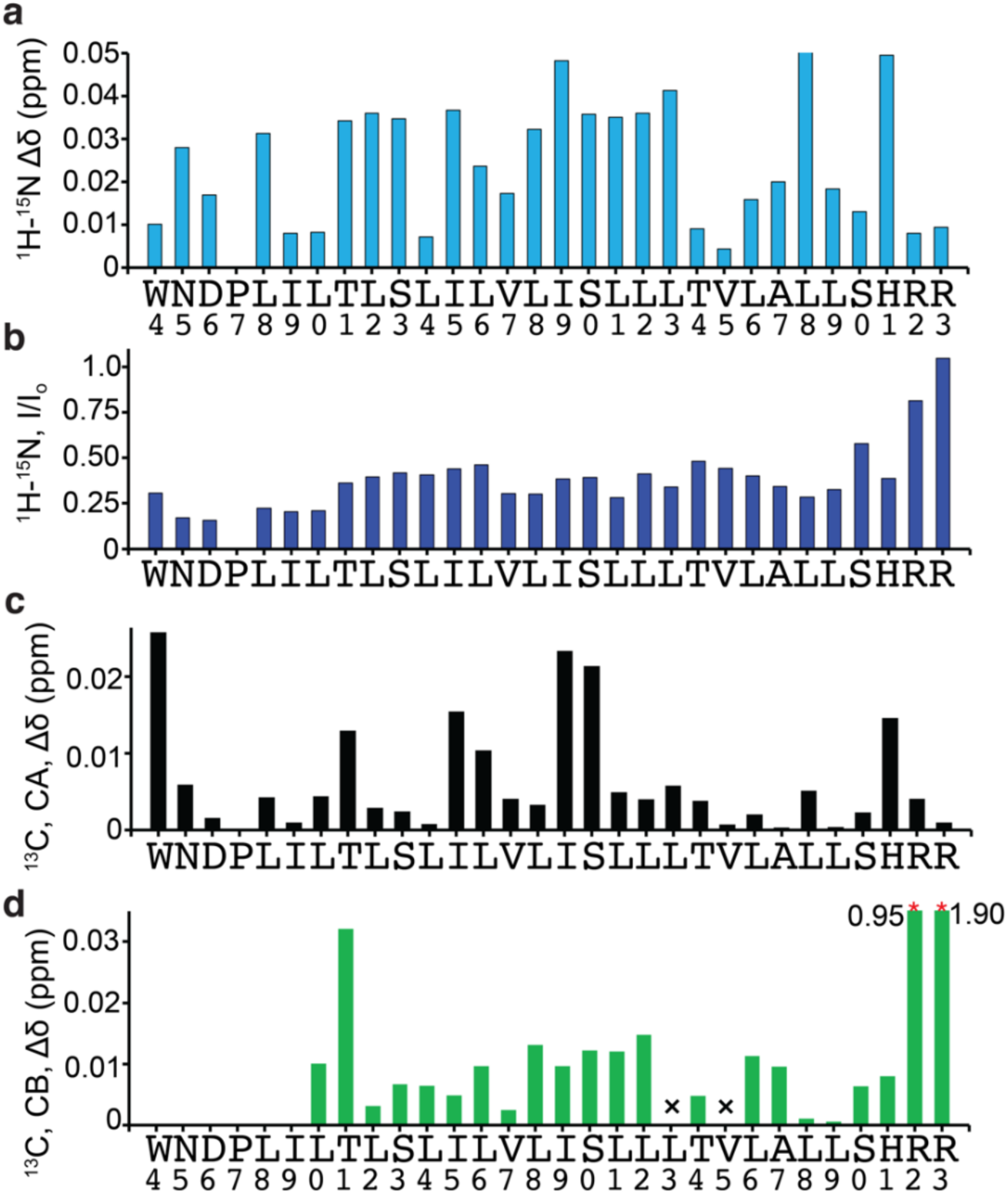
Per-residue chemical shift and intensity perturbations of mEpoR CHAMP-1 bound state versus monomeric state. (a) Hybrid ^1^H-^15^N chemical shift perturbation of monomeric and CHAMP-1 bound mEpoR-TM2. (b) Relative intensity (I) between ^1^H-^15^N resonances in HSQC spectra of monomeric and CHAMP-1 bound mEpoR- TM2. (c) CA chemical shift perturbation of monomeric and CHAMP-1 bound mEpoR-TM2. (d) CB chemical shift perturbation of monomeric and CHAMP-1 bound mEpoR-TM2. Missing residues notes by ‘x’. R32 and R33 are off-scale, and their induced shift values listed on top.

**Table S1.**
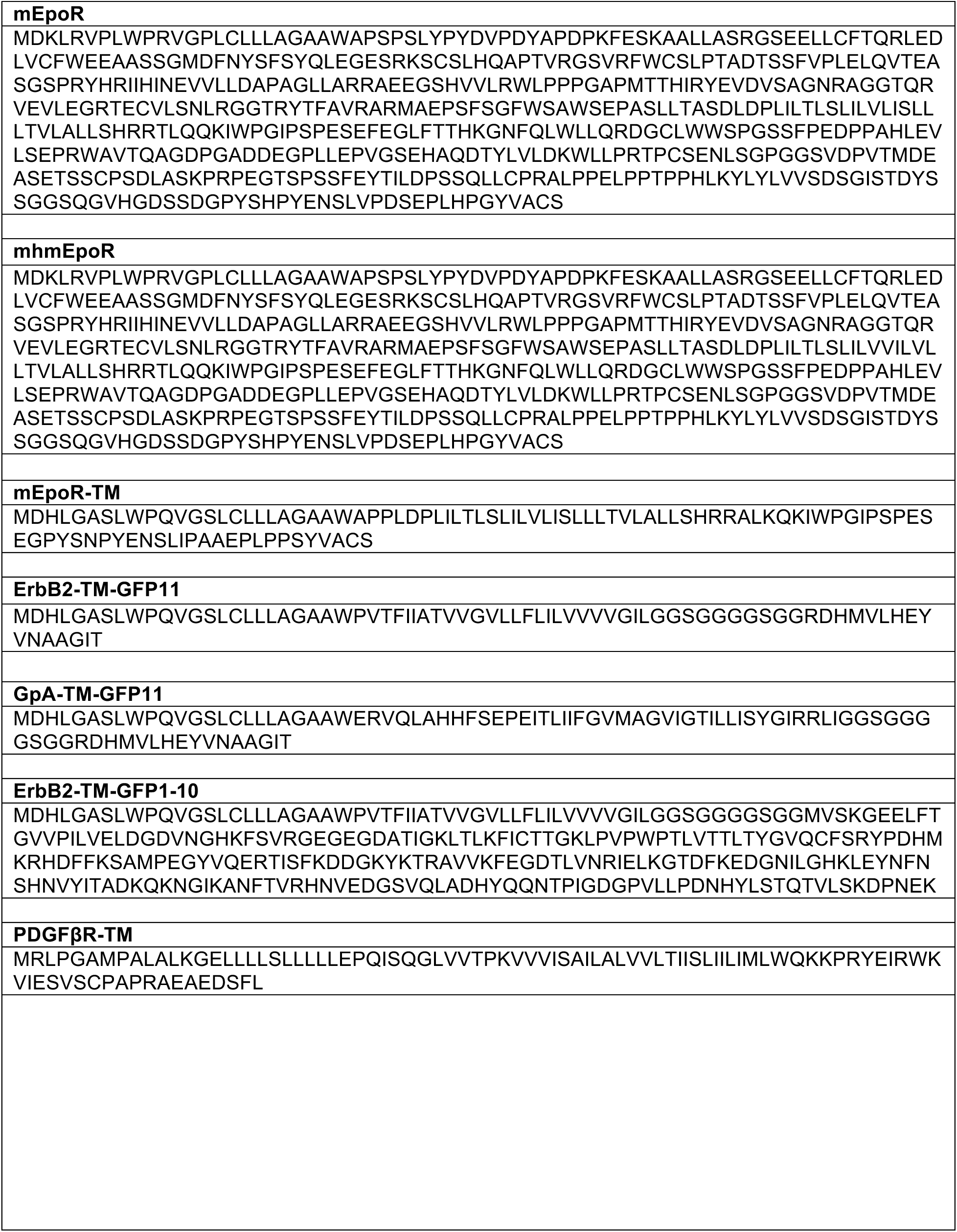
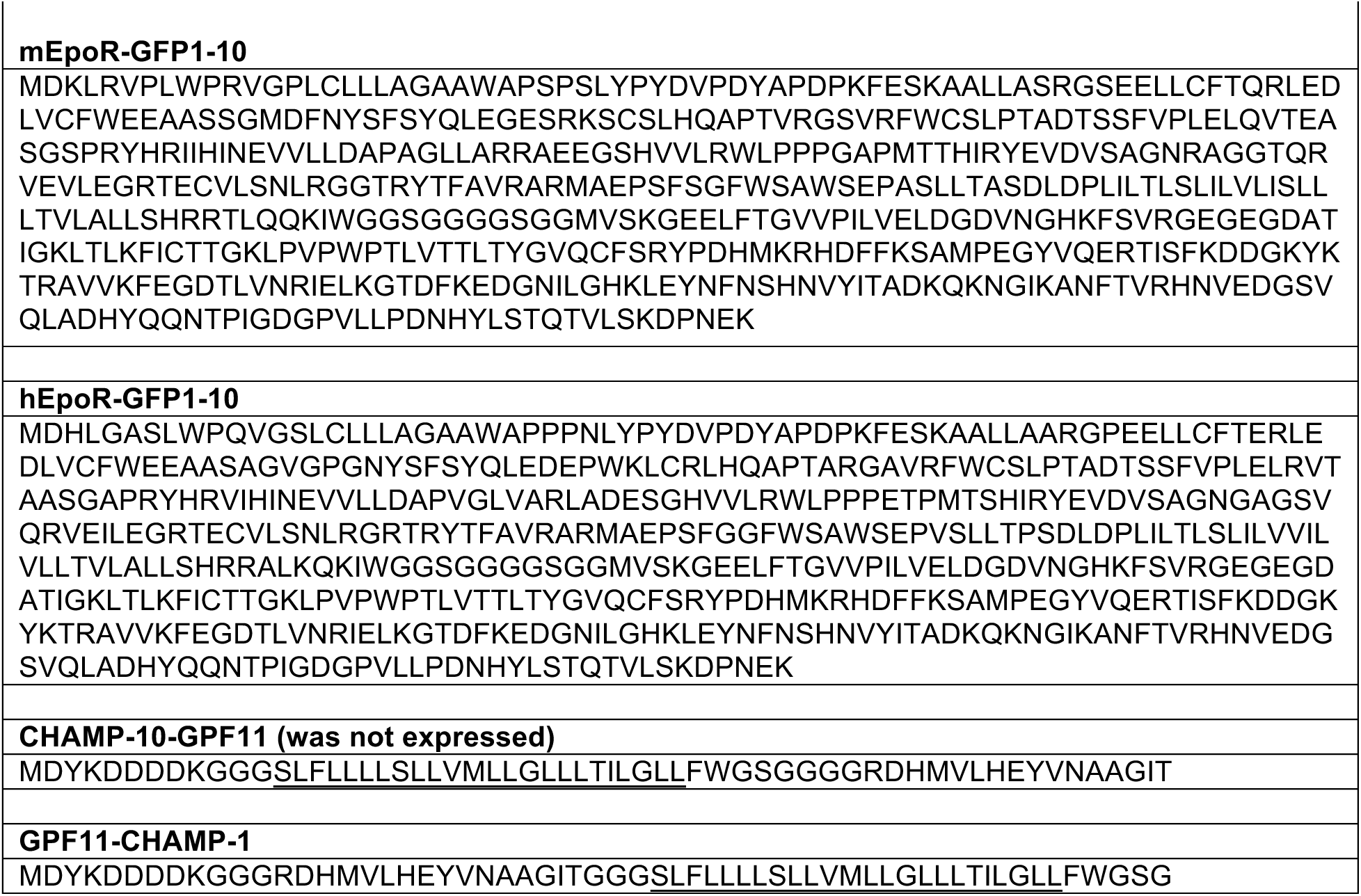
Expressed protein constructs in BaF3 cells. The designed CHAMP-1 TM domain underlined.

**Table S2.**
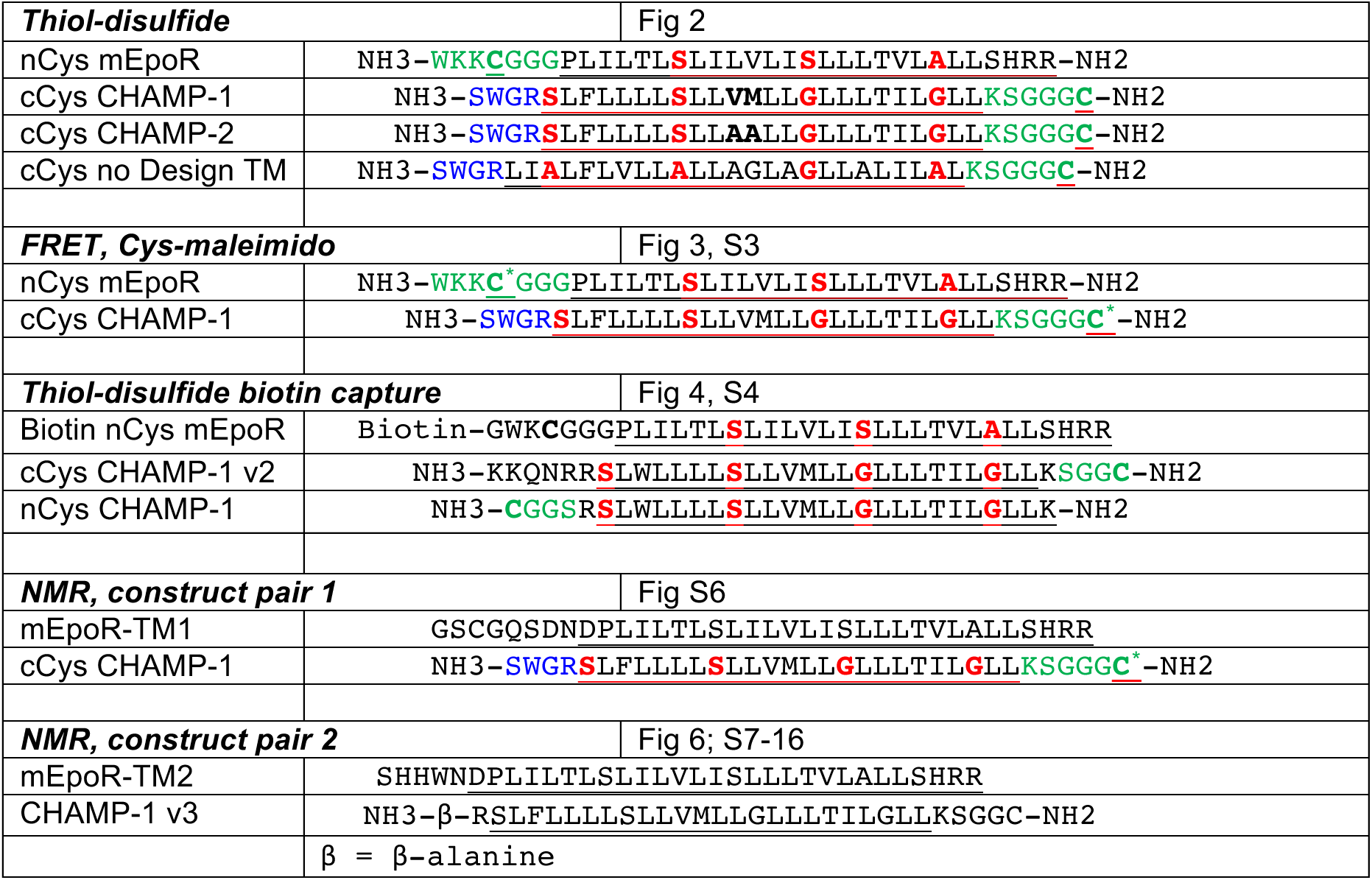
TM peptide sequences used in this work. Synthesized peptides were produced by solid-phase fmoc synthesis as C-terminal carboxamides and N-terminal free amines. “Biotin nCys mEpoR” was produced with biotin conjugated by peptide bond to the amino terminus. Fluorophore labeled TM peptides for FRET were produced as cysteine-maleimido conjugates at residues designated by asterisks. mEpoR-TM constructs for NMR were recombinantly expressed as fusion proteins: mEpoR-TM1 fused to T4 Lysozyme and cleaved by thrombin enzyme; mEpoR-TM2 fused to SUMO and cleaved chemically by Ni^2+^-induced SNAC self-cleavage.

**Table S3.**
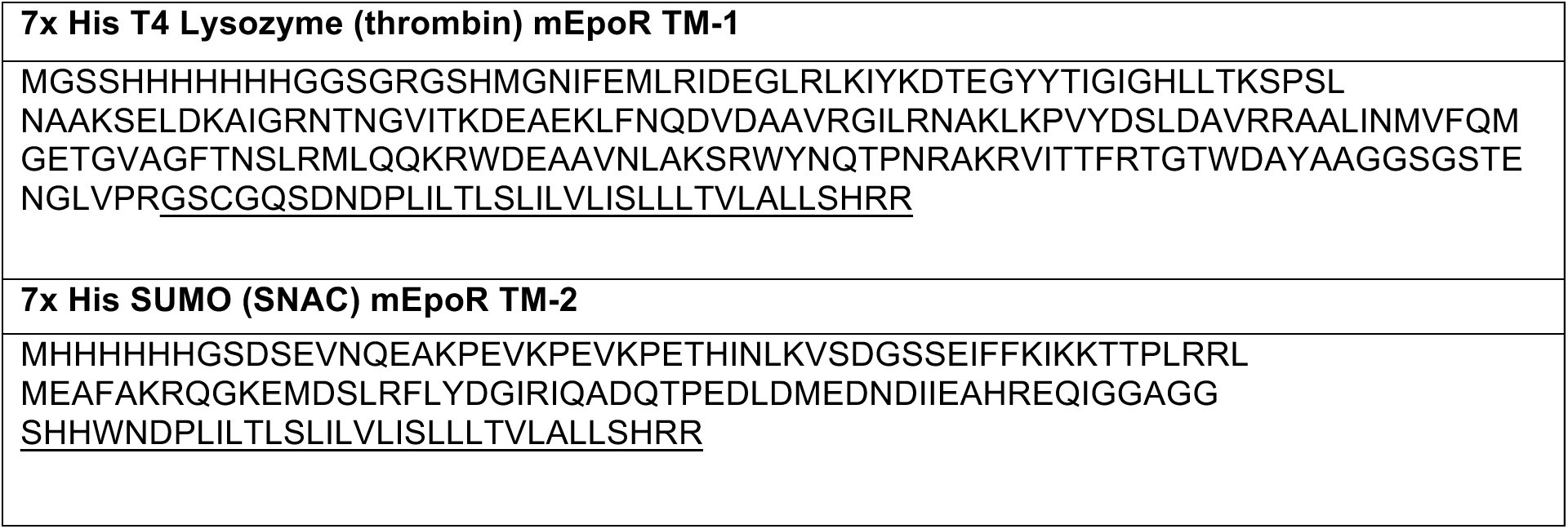
Fusion protein constructs for NMR. Constructs for isotopically enriched recombinant expression in *E. coli*. Final TM constructs after thrombin protease cleavage or sequence-specific nickel-assisted cleavage (SNAC) used for NMR are underlined.

**Table S4.** CB chemical shift perturbations. Induced chemical shift changes of mEpoR-TM in ^2^H-DPC upon addition of unlabeled CHAMP-1 peptide at 7 mol eqv. and 600 mol eqv. DPC, across multiple spectra including polypeptide carbon beta (CB) atoms resonances: HNCB, CBCANH, CCcoNH, ^1^H-^13^C HSQC. Frequencies listed in ppm. Complete resonance list for the monomeric mEpoR-TM in DPC can be found in the BMRB repository.

| Monomer mEpoR-TM, HNCB |  |  |  | Complex, CBCANH |  |  |  |  |
| --- | --- | --- | --- | --- | --- | --- | --- | --- |
|  | C | N | H |  | C | N | H |  |
| | w1 | w2 | w3 | | w1 | w2 | w3 | $\delta$ Shift |
| I9CB-N-H | 36.40 | 118.23 | 7.40 |  |  |  |  |  |
| L10CB-N-H | 42.14 | 122.01 | 8.43 | L10CB-T11N-H | 42.04 | 114.89 | 7.95 | 0.10 |
| T11CB-N-H | 68.50 | 114.75 | 7.92 | T11CB-L12N-H | 68.18 | 119.57 | 8.22 | 0.32 |
| L12CB-N-H | 41.86 | 119.50 | 8.20 | L12CB-S13N-H | 41.82 | 114.64 | 8.27 | 0.03 |
| S13CB-N-H | 62.94 | 114.66 | 8.31 |  |  |  |  |  |
| L14CB-N-H | 41.61 | 121.30 | 7.98 | L14CB-I15N-H | 41.64 | 118.34 | 8.11 | 0.03 |
| I15CB-N-H | 37.33 | 118.24 | 8.07 |  |  |  |  |  |
| L16CB-N-H | 41.47 | 119.27 | 8.28 | L16CB-V17N-H | 41.39 | 120.92 | 8.35 | 0.09 |
| V17CB-N-H | 31.30 | 121.10 | 8.37 | V17CB-L18N-H | 31.32 | 120.05 | 8.44 | 0.03 |
| L18CB-N-H | 41.69 | 119.97 | 8.41 | L18CB-I19N-H | 41.82 | 117.78 | 8.70 | 0.13 |
| I19CB-N-H | 37.56 | 117.28 | 8.69 | I19CB-S20N-H | 37.65 | 116.61 | 8.15 | 0.10 |
| S20CB-N-H | 62.67 | 116.76 | 8.19 | S20CB-L21N-H | 62.79 | 124.29 | 8.52 | 0.12 |
| L21CB-N-H | 42.01 | 124.30 | 8.49 |  |  |  |  |  |
| L22CB-N-H | 41.79 | 119.51 | 8.33 |  |  |  |  |  |
| L23CB-N-H | 41.75 | 117.08 | 8.60 |  |  |  |  |  |
| T24CB-N-H | 67.80 | 116.15 | 7.90 | T24CB-V25N-H | 67.85 | 120.46 | 8.28 | 0.05 |
| V25CB-N-H | 31.35 | 120.47 | 8.28 |  |  |  |  |  |
| L26CB-N-H | 41.58 | 118.71 | 8.36 | L26CB-A27N-H | 41.61 | 121.88 | 8.49 | 0.02 |
| A27CB-N-H | 18.32 | 121.93 | 8.53 | A27CB-L28N-H | 18.42 | 118.39 | 8.31 | 0.10 |
| L28CB-N-H | 42.26 | 118.43 | 8.35 | L28CB-L29N-H | 42.26 | 117.85 | 8.44 | 0.01 |
| L29CB-N-H | 42.24 | 117.89 | 8.46 |  |  |  |  |  |
| S30CB-N-H | 63.83 | 112.87 | 7.91 | S30CB-H31N-H | 63.89 | 118.29 | 7.83 | 0.05 |
| H31CB-N-H | 28.92 | 118.14 | 7.87 |  |  |  |  |  |
| R32CB-N-H | 30.85 | 126.07 | 7.76 |  |  |  |  |  |
| R33CB-N-H | 31.76 | 126.08 | 7.75 |  |  |  |  |  |

**Table S4.**
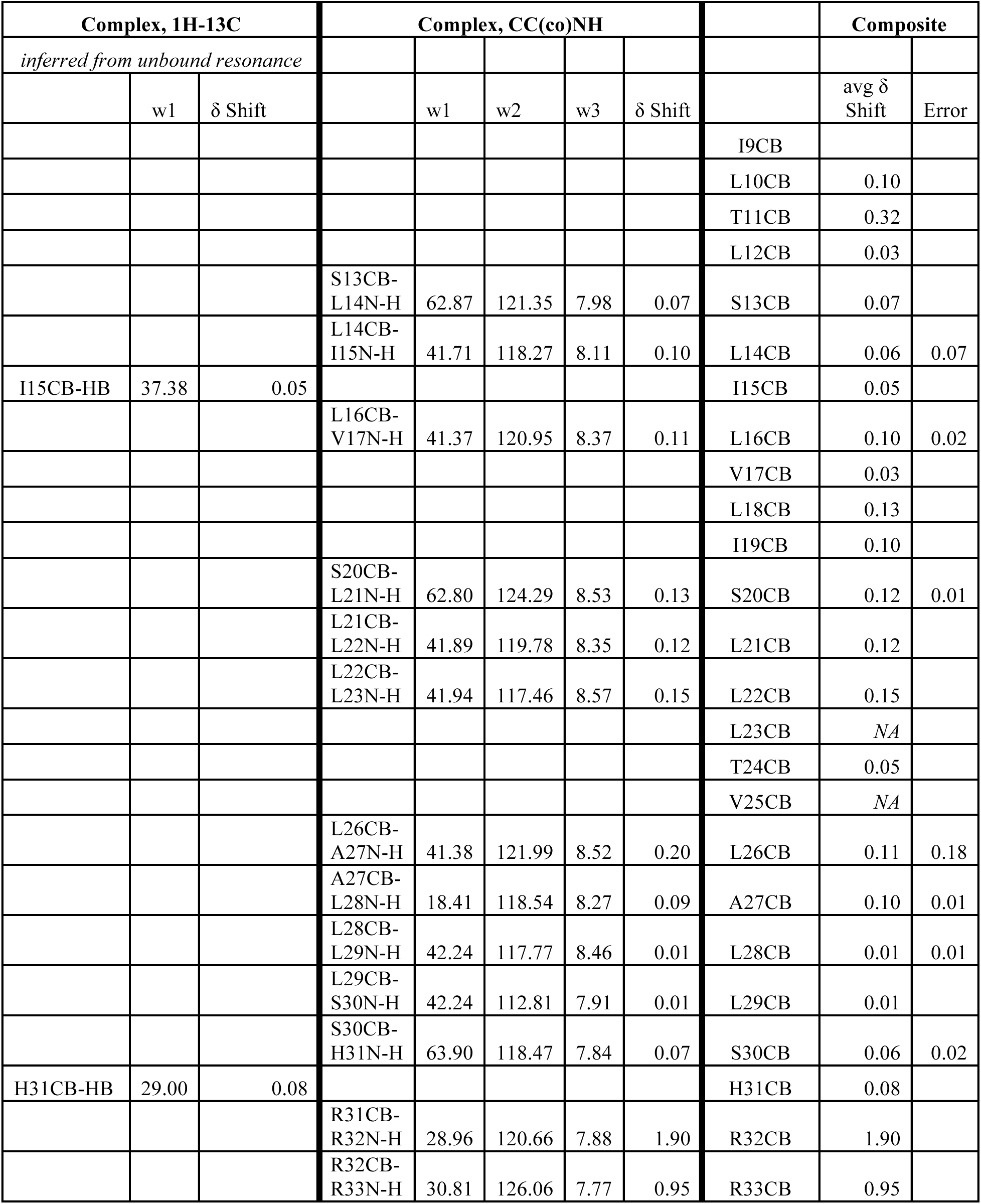
CB chemical shift perturbations, continued. Induced chemical shift changes of mEpoR-TM in ^2^H-DPC upon addition of unlabeled CHAMP-1 peptide at 7 mol eqv. and 600 mol eqv. DPC, across multiple spectra including polypeptide carbon beta (CB) atoms resonances: HNCB, CBCANH, CCcoNH, ^1^H-^13^C HSQC. Frequencies listed in ppm.

**Table S5.**
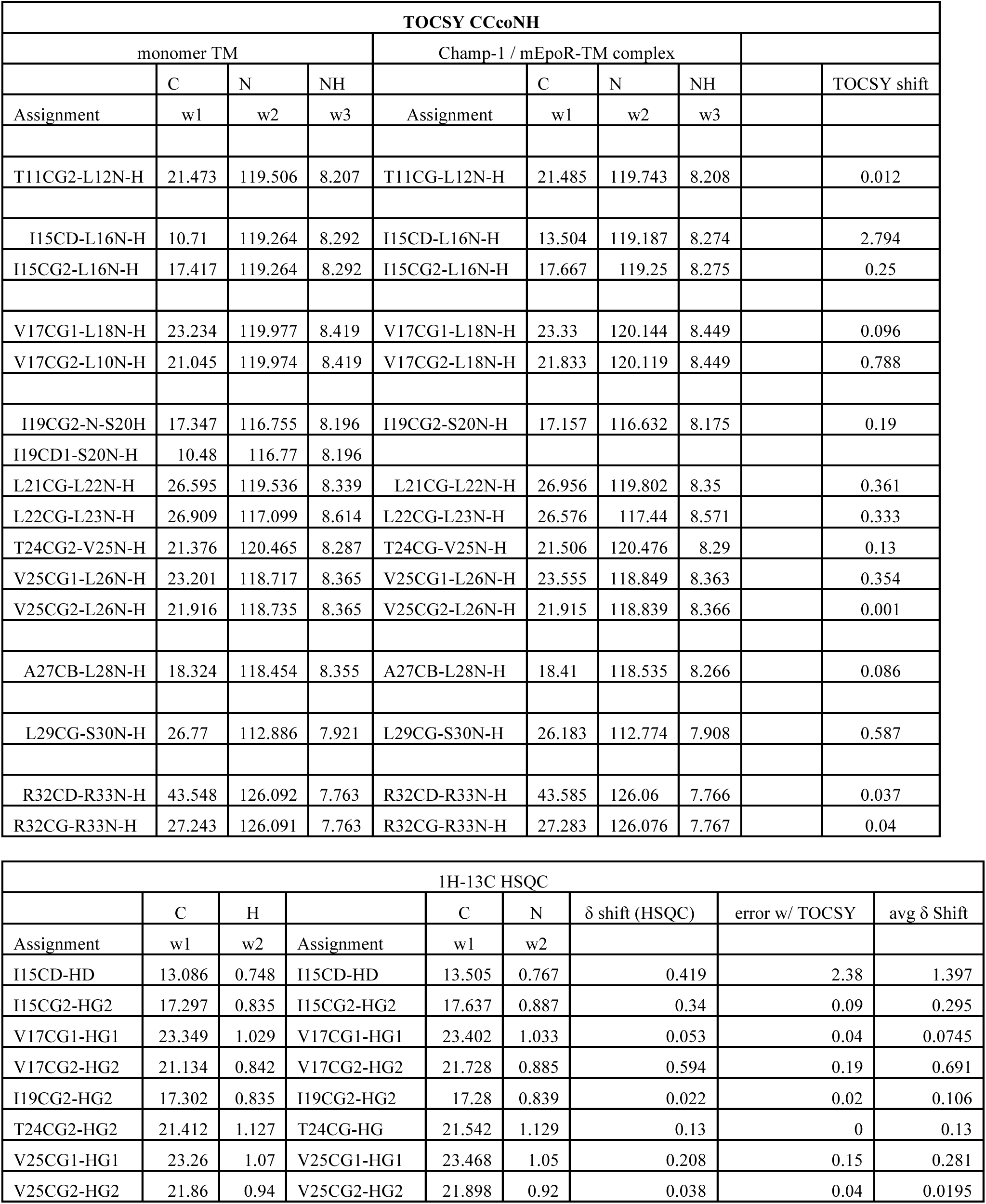
^13^C Sidechain chemical shift perturbations. Detectable and unambiguous shift changes of mEpoR-TM in ^2^H-DPC with unlabeled CHAMP-1 in CCcoNH and ^1^H-^13^C HSQC spectra beyond CB. Frequencies listed in ppm.

| TOCSY CCcoNH |  |  |  |  |  |  |  |  |  |
| --- | --- | --- | --- | --- | --- | --- | --- | --- | --- |
| monomer TM |  |  |  | Champ-1 / mEpoR-TM complex |  |  |  |  |  |
|  | C | N | NH |  | C | N | NH |  | TOCSY shift |
| Assignment | w1 | w2 | w3 | Assignment | w1 | w2 | w3 |  |  |
| T11CG2-L12N-H | 21.473 | 119.506 | 8.207 | T11CG-L12N-H | 21.485 | 119.743 | 8.208 |  | 0.012 |
| I15CD-L16N-H | 10.71 | 119.264 | 8.292 | I15CD-L16N-H | 13.504 | 119.187 | 8.274 |  | 2.794 |
| I15CG2-L16N-H | 17.417 | 119.264 | 8.292 | I15CG2-L16N-H | 17.667 | 119.25 | 8.275 |  | 0.25 |
| V17CG1-L18N-H | 23.234 | 119.977 | 8.419 | V17CG1-L18N-H | 23.33 | 120.144 | 8.449 |  | 0.096 |
| V17CG2-L10N-H | 21.045 | 119.974 | 8.419 | V17CG2-L18N-H | 21.833 | 120.119 | 8.449 |  | 0.788 |
| I19CG2-N-S20H | 17.347 | 116.755 | 8.196 | I19CG2-S20N-H | 17.157 | 116.632 | 8.175 |  | 0.19 |
| I19CD1-S20N-H | 10.48 | 116.77 | 8.196 |  |  |  |  |  |  |
| L21CG-L22N-H | 26.595 | 119.536 | 8.339 | L21CG-L22N-H | 26.956 | 119.802 | 8.35 |  | 0.361 |
| L22CG-L23N-H | 26.909 | 117.099 | 8.614 | L22CG-L23N-H | 26.576 | 117.44 | 8.571 |  | 0.333 |
| T24CG2-V25N-H | 21.376 | 120.465 | 8.287 | T24CG-V25N-H | 21.506 | 120.476 | 8.29 |  | 0.13 |
| V25CG1-L26N-H | 23.201 | 118.717 | 8.365 | V25CG1-L26N-H | 23.555 | 118.849 | 8.363 |  | 0.354 |
| V25CG2-L26N-H | 21.916 | 118.735 | 8.365 | V25CG2-L26N-H | 21.915 | 118.839 | 8.366 |  | 0.001 |
| A27CB-L28N-H | 18.324 | 118.454 | 8.355 | A27CB-L28N-H | 18.41 | 118.535 | 8.266 |  | 0.086 |
| L29CG-S30N-H | 26.77 | 112.886 | 7.921 | L29CG-S30N-H | 26.183 | 112.774 | 7.908 |  | 0.587 |
| R32CD-R33N-H | 43.548 | 126.092 | 7.763 | R32CD-R33N-H | 43.585 | 126.06 | 7.766 |  | 0.037 |
| R32CG-R33N-H | 27.243 | 126.091 | 7.763 | R32CG-R33N-H | 27.283 | 126.076 | 7.767 |  | 0.04 |

| $^1\text{H}$-$^{13}\text{C}$ HSQC | | | | | | | | |
| --- | --- | --- | --- | --- | --- | --- | --- | --- |
| | C | H | | C | N | $\delta$ shift (HSQC) | error w/ TOCSY | avg $\delta$ Shift |
| Assignment | w1 | w2 | Assignment | w1 | w2 |  |  |  |
| I15CD-HD | 13.086 | 0.748 | I15CD-HD | 13.505 | 0.767 | 0.419 | 2.38 | 1.397 |
| I15CG2-HG2 | 17.297 | 0.835 | I15CG2-HG2 | 17.637 | 0.887 | 0.34 | 0.09 | 0.295 |
| V17CG1-HG1 | 23.349 | 1.029 | V17CG1-HG1 | 23.402 | 1.033 | 0.053 | 0.04 | 0.0745 |
| V17CG2-HG2 | 21.134 | 0.842 | V17CG2-HG2 | 21.728 | 0.885 | 0.594 | 0.19 | 0.691 |
| I19CG2-HG2 | 17.302 | 0.835 | I19CG2-HG2 | 17.28 | 0.839 | 0.022 | 0.02 | 0.106 |
| T24CG2-HG2 | 21.412 | 1.127 | T24CG-HG | 21.542 | 1.129 | 0.13 | 0 | 0.13 |
| V25CG1-HG1 | 23.26 | 1.07 | V25CG1-HG1 | 23.468 | 1.05 | 0.208 | 0.15 | 0.281 |
| V25CG2-HG2 | 21.86 | 0.94 | V25CG2-HG2 | 21.898 | 0.92 | 0.038 | 0.04 | 0.0195 |

